# Hemispheric Asymmetry of Tau Pathology is Related to Asymmetric Amyloid Deposition in Alzheimer’s Disease

**DOI:** 10.1101/2025.04.15.648728

**Authors:** T.E. Anijärv, R. Ossenkoppele, R. Smith, A. P. Binette, L.E. Collij, H.H. Behjat, J. Rittmo, L. Karlsson, K. Ahmadi, O. Strandberg, the Alzheimer’s Disease Neuroimaging Initiative, D. van Westen, J.W. Vogel, E. Stomrud, S. Palmqvist, N. Mattsson-Carlgren, N. Spotorno, O. Hansson

**Affiliations:** Clinical Memory Research Unit, Department of Clinical Sciences Malmö, Faculty of Medicine, Lund University, Lund, Sweden; Alzheimer Center Amsterdam, Neurology, Vrije Universiteit Amsterdam, Amsterdam UMC, Amsterdam, the Netherlands; Neurodegeneration, Amsterdam Neuroscience, Amsterdam, the Netherlands; Memory Clinic, Skåne University Hospital, Malmö, Sweden; Department of Physiology and Pharmacology, Université de Montréal, Montréal, Quebec, Canada; Centre de Recherche de l’Institut Universitaire de Gériatrie de Montréal, Montréal, Quebec, Canada; Radiology and Nuclear Medicine, Vrije Universiteit Amsterdam, Amsterdam UMC, Amsterdam, the Netherlands; Brain Imaging, Amsterdam Neuroscience, Amsterdam, the Netherlands; SciLifeLab, Department of Clinical Sciences Malmö, Faculty of Medicine, Lund University, Lund, Sweden; Department of Neuropsychology, Ruhr University Bochum, Bochum, Germany; Diagnostic Radiology, Institution for Clinical Sciences, Lund University, Lund, Sweden; Wallenberg Center for Molecular Medicine, Lund University, Lund, Sweden

## Abstract

The distribution of tau pathology in Alzheimer’s disease (AD) shows remarkable inter-individual heterogeneity, including hemispheric asymmetry. However, the factors driving this asymmetry remain poorly understood. We explored whether tau asymmetry is linked to i) reduced inter-hemispheric brain connectivity (potentially restricting tau spread), or ii) asymmetry in amyloid-beta (Aβ) distribution (indicating greater hemisphere-specific vulnerability to AD pathology). 452 participants from the Swedish BioFINDER-2 cohort with evidence of both Aβ pathology (CSF Aβ42/40 or neocortical Aβ-PET) and tau pathology (temporal tau-PET), were categorised as left asymmetric (n=102), symmetric (n=306), or right asymmetric (n=44) based on temporal lobe tau-PET uptake distribution. Edge-wise inter-hemispheric functional (RSfMRI; n=318) and structural connectivity (dMRI; n=352) patterns were examined but no differences in inter-hemispheric functional or structural connectivity were found between groups. However, a strong association was observed between tau and Aβ laterality patterns based on PET uptake (n=233; β=0.632, p<0.001), which was replicated in three independent cohorts (n=234; β=0.535, p<0.001). In a longitudinal Aβ-positive sample, baseline Aβ asymmetry predicted the progression of tau laterality over time (n=289; β=0.025, p=0.028). These findings suggest that tau asymmetry is not associated with a weaker inter-hemispheric connectivity but might reflect hemispheric differences in vulnerability to Aβ pathology, underscoring the role of regional vulnerability in determining the distribution of AD pathology.

## Introduction

Alzheimer’s disease (AD), the leading cause of dementia worldwide, exhibits substantial heterogeneity in its pathological manifestations and clinical progression.^1,2^ The disease is characterized by two primary pathological hallmarks: the accumulation of amyloid-beta (Aβ) plaques throughout the neocortex and the progressive aggregation of hyperphosphorylated tau proteins forming neurofibrillary tangles, ultimately resulting in neurodegeneration and cognitive decline.^3^ The accumulation of tau pathology in AD is commonly described as following a stereotypical distribution.^4,5^ Nonetheless, tau-PET studies have demonstrated heterogeneity in the distribution of tau across individuals and multiple spatiotemporal patterns have been described, differentially associated with cognitive functioning and decline.^1,2,6–8^

One of the manifestations of this heterogeneity is an asymmetric distribution of tau pathology between the two hemispheres. Hemispheric asymmetry of tau pathology in AD has been associated with younger age, more severe pathological burden and rapid multi-domain cognitive impairment.^7,9–11^ Moreover, some studies indicate that tau asymmetry is more easily identified during the more advanced stages of the disease,^7,12^ often with left hemisphere dominance.^13^ However, asymmetrical tau distribution has also been found to be present in preclinical AD.^10^ More commonly, asymmetry in tau distribution is presented in atypical AD cases, such as posterior cortical atrophy (PCA)^14^ and primary progressive aphasia (PPA)^15^ where PCA can present with both left and right predominant tau distribution while PPA is usually characterized by left-sided asymmetry.^16–18^ Furthermore, a data-driven approach to tau-PET data in a large multi-cohort study^7^, uncovered four primary AD subtypes characterized by different spatiotemporal profiles of tau pathology. Among these, the lateral temporal subtype, comprising 19% of cases, exhibited strong tau asymmetry, while the remaining subtypes demonstrated more moderate patterns of tau lateralisation.

Although previous work indicates that asymmetric distribution of tau pathology is relatively common, the underlying drivers of this phenomenon are not clearly characterised. Two primary mechanisms have been proposed to explain tau accumulation patterns. The first suggests that misfolded tau proteins spread through connected brain regions in a prion-like manner.^19–23^ The second mechanism proposes that tau accumulation is primarily driven by local replication or regional vulnerability, which can be influenced by various factors including the local presence of Aβ pathology.^24–26^ These mechanisms are not mutually exclusive. Understanding their contribution to hemispheric tau asymmetry could reveal critical insights into AD pathophysiology and guide the development of novel therapeutic strategies. In this study, we proposed two possible hypotheses for explaining asymmetric tau distribution. First, we investigated whether individuals characterized by asymmetric tau patterns display reduced inter-hemispheric functional and structural connectivity, possibly indicating reduced tau spreading between hemispheres. Second, we explored the idea that this asymmetry in tau accumulation is related to hemispheric differences in Aβ deposition, suggesting that these individuals display greater regional vulnerability to AD pathology in one hemisphere more than the other.

## Results

### Participants

A sample of 837 Aβ-positive (A+) participants, based on neocortical Aβ-PET or CSF Aβ42/40, with available tau-PET scan(s) from the BioFINDER-2 cohort was included in this study. The cohort consisted of both clinically unimpaired and clinically impaired participants (see ‘Methods – Participants’ for detailed information). A cross-sectional sample of subjects with evidence of tau pathology (A+T+; n=475) based on unilateral (i.e., most affected hemisphere) tau-PET uptake in the temporal meta region of interest (meta-ROI; SUVR>1.362^27,28^) was selected and categorized into three groups according to the spatial distribution of tau. Specifically, a tau laterality index (LI) was computed for each participant based on the tau-PET uptake in a temporal meta-ROI covering the regions described by Braak stages I-IV. For the following analyses, tau laterality was examined both as a continuous measure (tau LI) and as a grouping factor. Three groups were defined: participants with LI exceeding ±1 SD from perfect symmetry (i.e., LI=0) were assigned to the right asymmetric (RA; n=44) or left asymmetric (LA; n=102) group, respectively, while those within ±1 SD were assigned to the symmetric (S; n=306) group. Subjects with a borderline laterality index, defined as a ±5% interval around the threshold were not allocated to any group and excluded from the analyses (n=23), resulting in a final sample size of 452 (Table 1; Fig. 1ab).

**Figure 1.**
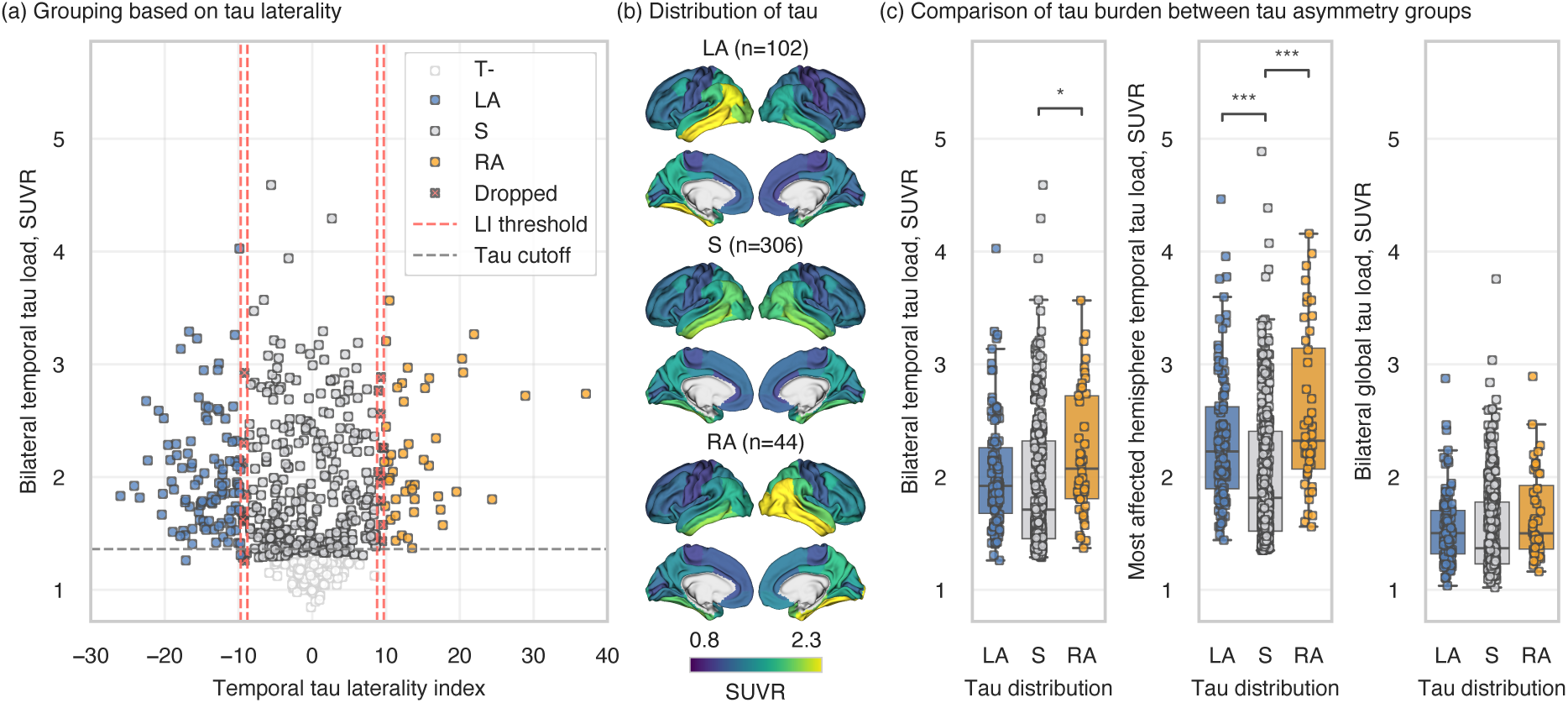
Grouping of the subjects: (a) Participants were divided into three groups based on the distribution of temporal tau-PET uptake; (b) Average tau-PET SUVRs for each group; (c) Comparison of tau load between the groups. Laterality index (%) = 100 × (right tau – left tau) / (right tau + left tau). LA – left tau asymmetric; S – tau symmetric – RA – right tau asymmetric; SUVR – standardised uptake value ratio; * – p<0.05; *** – p<0.001.

**Table 1.**
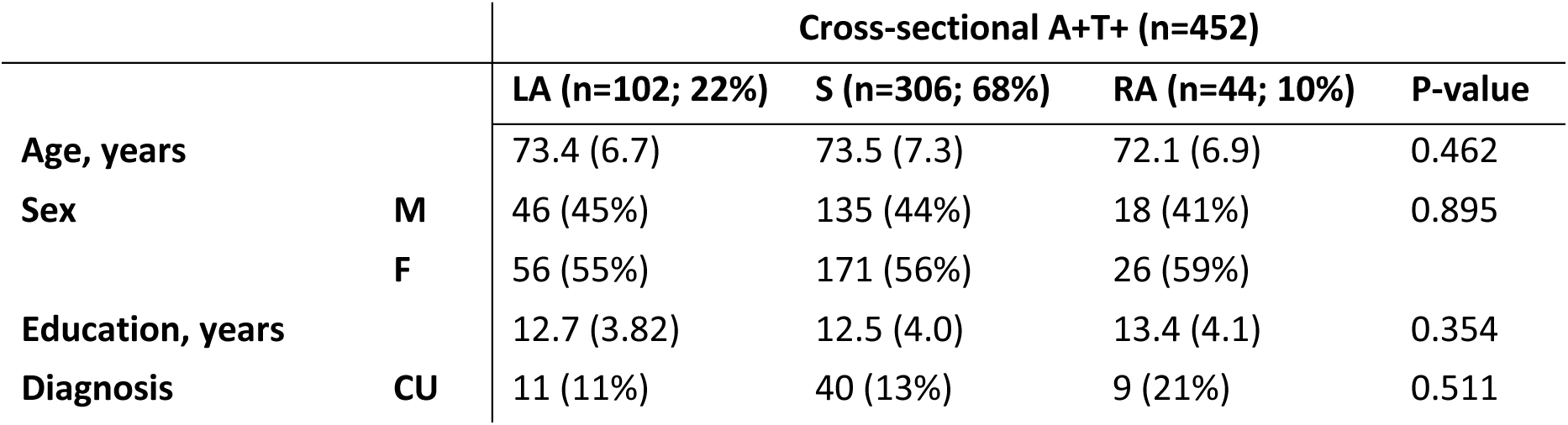
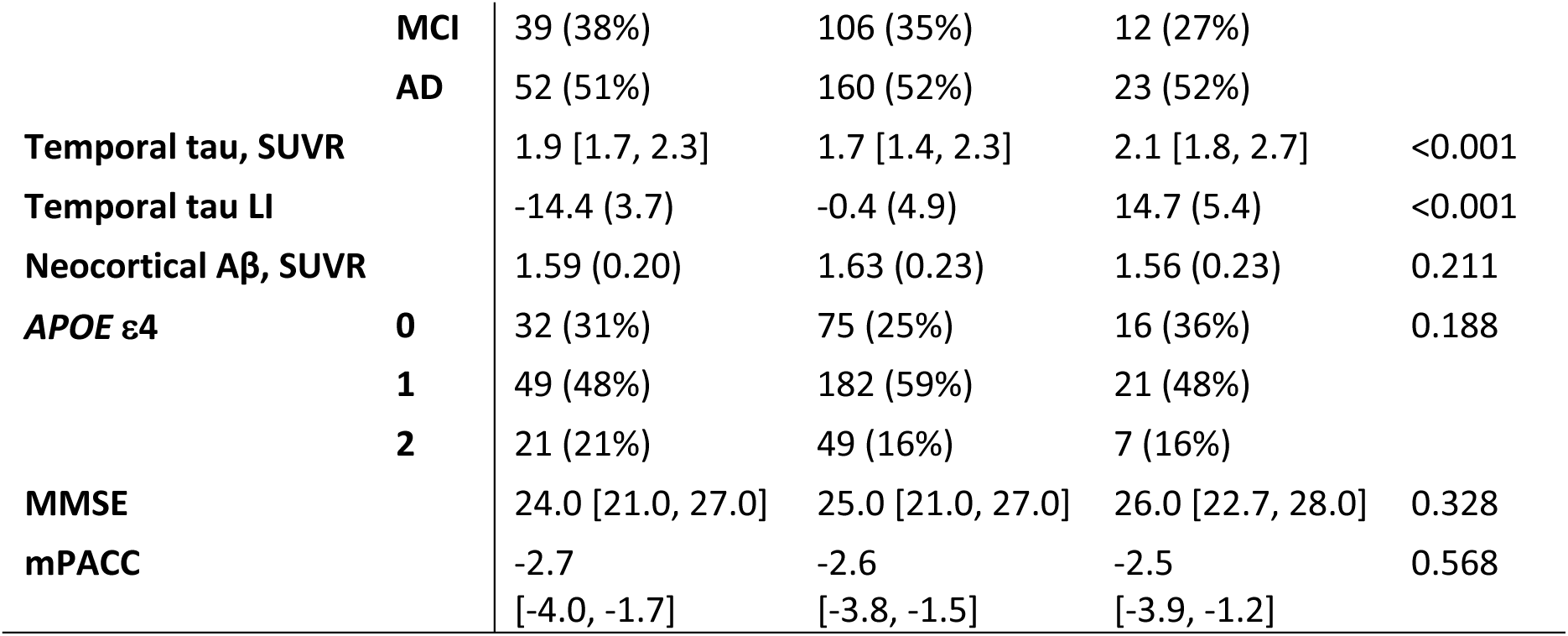
Demographics. Categorical variables have been presented as ’count (%)’, normally distributed continuous variables as ’mean (SD)’ and non-normally distributed variables as ‘median [IQR]’. LA – left tau asymmetric; S – tau symmetric – RA – right tau asymmetric; M – male; F – female; CU – cognitively unimpaired; MCI – mild cognitive impairment; AD – Alzheimer’s disease; SUVR – standardised uptake value ratio; LI – laterality index; Aβ – amyloid-beta; MMSE – Mini-Mental State Examination; mPACC – modified Preclinical Alzheimer Cognitive Composite.

|  |  | Cross-sectional A+T+ (n=452) |  |  |  |
| --- | --- | --- | --- | --- | --- |
|  |  | LA (n=102; 22%) | S (n=306; 68%) | RA (n=44; 10%) | P-value |
| Age, years |  | 73.4 (6.7) | 73.5 (7.3) | 72.1 (6.9) | 0.462 |
| Sex | M | 46 (45%) | 135 (44%) | 18 (41%) | 0.895 |
|  | F | 56 (55%) | 171 (56%) | 26 (59%) |  |
| Education, years |  | 12.7 (3.82) | 12.5 (4.0) | 13.4 (4.1) | 0.354 |
| Diagnosis | CU | 11 (11%) | 40 (13%) | 9 (21%) | 0.511 |
|  | **MCI** | 39 (38%) | 106 (35%) | 12 (27%) |  |
|  | **AD** | 52 (51%) | 160 (52%) | 23 (52%) |  |
| **Temporal tau, SUVR** |  | 1.9 [1.7, 2.3] | 1.7 [1.4, 2.3] | 2.1 [1.8, 2.7] | <0.001 |
| **Temporal tau LI** |  | -14.4 (3.7) | -0.4 (4.9) | 14.7 (5.4) | <0.001 |
| **Neocortical Aβ, SUVR** |  | 1.59 (0.20) | 1.63 (0.23) | 1.56 (0.23) | 0.211 |
| **APOE ε4** | **0** | 32 (31%) | 75 (25%) | 16 (36%) | 0.188 |
|  | **1** | 49 (48%) | 182 (59%) | 21 (48%) |  |
|  | **2** | 21 (21%) | 49 (16%) | 7 (16%) |  |
| **MMSE** |  | 24.0 [21.0, 27.0] | 25.0 [21.0, 27.0] | 26.0 [22.7, 28.0] | 0.328 |
| **mPACC** |  | -2.7<br>[-4.0, -1.7] | -2.6<br>[-3.8, -1.5] | -2.5<br>[-3.9, -1.2] | 0.568 |

Demographic characteristics, including age, sex, *APOE* ε4 allele carriership, and cognitive performance did not differ between the three groups. Moreover, average bilateral global (i.e., both whole hemispheres) tau load did not differ between the three groups, nor did Aβ (Table 1; Fig. 1c). SUVR values from the temporal meta-ROI showed a trend toward higher average bilateral tau load in participants with asymmetric tau distribution compared to those with symmetric distribution, though this difference was statistically significant only between the right tau asymmetric group and the symmetric group (t=2.688, p=0.023). When examining unilateral tau load in the temporal meta-ROI in the most affected hemisphere, both asymmetric groups showed significantly higher load compared to the symmetric group (LA-S: t=4.828, p<0.001; RA-S: t=5.240, p<0.001). To account for these differences, SUVR values from average bilateral global tau load were included as a covariate in subsequent analyses.

### Inter-hemispheric connectivity does not differ between participants with asymmetric and symmetric pattern of tau deposition

Subsequently, we assessed whether asymmetric tau distribution is associated with differences in inter-hemispheric brain connectivity. Connectivity was calculated between Desikan-Killiany atlas regions using functional connectivity (correlation of blood-oxygen-level-dependent signals, BOLD) and structural connectivity (anatomically-constrained tractography). To ensure biological relevance, only the top 10% of inter-hemispheric connections identified in a separate sample of healthy controls were retained for analysis (see ‘Methods’ for detailed information). No statistically significant differences were found in average inter-hemispheric functional connectivity (n=318; S-LA: β=-0.151, 95%CI=[-0.417; 0.116], p=0.802; S-RA: β=0.076, 95%CI=[-0.312; 0.463], p>0.9) or structural connectivity (n=352; S-LA: β=0.135, 95%CI=[-0.107; 0.376], p=0.823; S-RA: β=0.170, 95%CI=[-0.162; 0.502], p>0.9) between the tau asymmetric groups and the tau symmetric group (Fig. 2a). Similarly, edge-wise comparisons of inter-hemispheric connections showed no significant differences between the groups. When examining connectivity between all pairs of contralateral regions (n=1412 connections), no differences were found between symmetric and asymmetric groups in either functional (all p>0.3 for S-LA and all p>0.2 for S-RA comparisons) or structural connectivity (all p>0.2 for S-LA and all p>0.4 for S-RA comparisons). To confirm that the results did not arise from the group definitions, we then further performed additional linear regression analysis using the continuous tau laterality index instead of group comparisons (see Supplementary Fig. S2.1). There was no significant association between global (i.e., average of all inter-hemispheric edges) tau laterality index and functional connectivity (β=0.090, 95%CI=[-0.027; 0.207], p=0.130) or structural connectivity (β=-0.037, 95%CI=[-0.144; 0.071], p=0.502).

**Figure 2.**
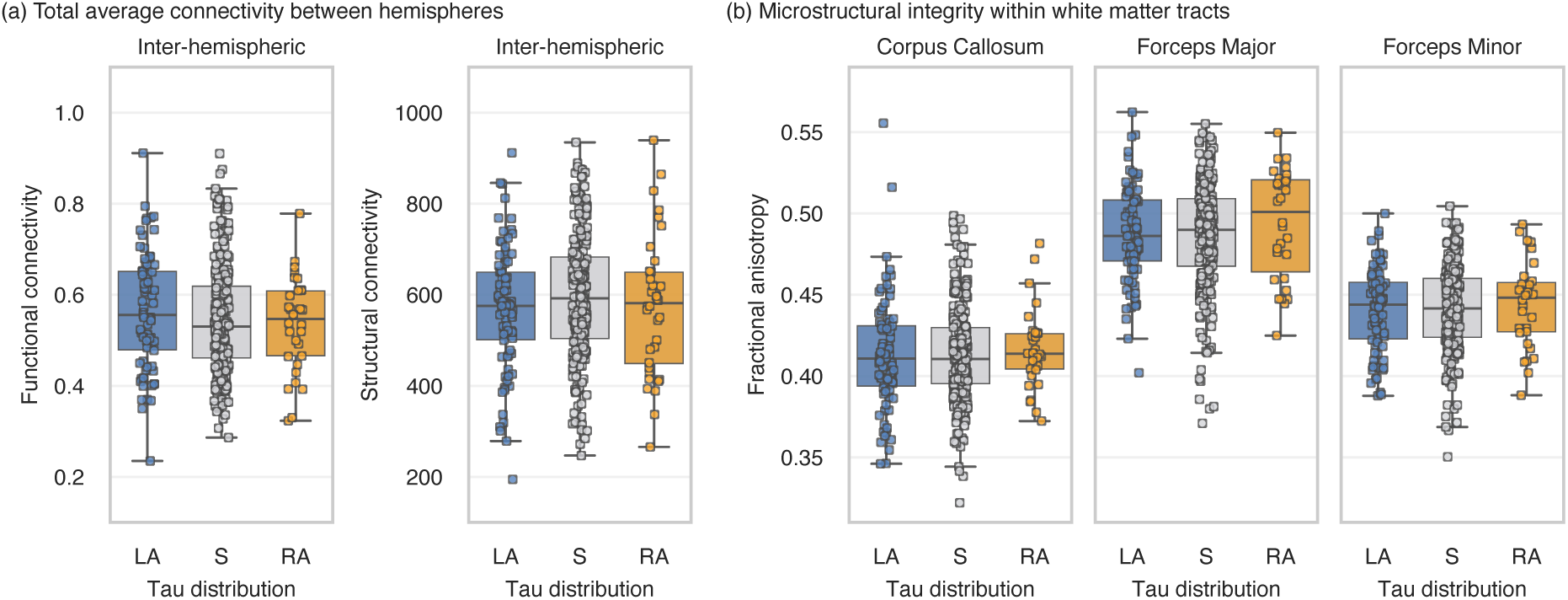
Comparison of average connectivity between the tau asymmetry groups: (a) Global functional and structural connectivity; (b) Fractional anisotropy in main white matter tracts connecting the two hemispheres. Note – 16 subjects were dropped from the latter analyses after visual quality control of the tract segmentation, resulting in n=336. LA – left tau asymmetric; S – tau symmetric – RA – right tau asymmetric.

Microstructural integrity was assessed within the main white matter tracts connecting the two hemispheres between the groups. There were no differences in fractional anisotropy in neither of the inter-hemispheric tracts (Fig. 2b) – in the corpus callosum (S-LA: β=0.042, 95%CI=[- 0.216; 0.300], p>0.9; S-RA: β=-0.134, 95%CI=[-0.518; 0.251], p>0.9), forceps major (S-LA: β=-0.033, 95%CI=[-0.289; 0.223], p>0.9; S-RA: β=-0.220, 95%CI=[-0.599; 0.159], p=0.762), or forceps minor (S-LA: β=0.001, 95%CI=[-0.258; 0.260], p>0.9; S-RA: β=-0.136, 95%CI=[-0.520; 0.248], p>0.9); similar results were found for mean diffusivity (see Supplementary Fig. S2.2).

Moreover, performing a whole-brain connectome analysis (i.e., not limited to only inter-hemispheric connections) using Network Based Statistics (NBS; see Supplementary Fig. S2.3 and Table S2.1) did not show evidence of reduced functional or structural connectivity between hemispheres when comparing the asymmetric and symmetric groups (see Supplementary S2 for a full overview of the NBS results).

### Strong associations between the laterality of Aβ and tau pathologies

We then assessed whether tau asymmetry is related to hemispheric differences in vulnerability to Aβ pathology. Within the A+T+ sample with available Aβ-PET (n=233), there was a strong association between the degree of global laterality in tau and in Aβ pathologies (Fig. 3a; β=0.632, 95%CI=[0.530; 0.733], p<0.001); however, the magnitude of asymmetry was greater in tau than in Aβ pathology. Similar results were found when investigating the associations at a regional level with all regions (min β=0.167; mean β=0.380; max β=0.633; all p<0.05) except for two regions (the rostral anterior cingulate [β=0.130, p=0.051] and the hippocampus [β=-0.034, p=0.608]). The strongest effect sizes were within the temporal regions, particularly the inferior temporal (β=0.633, p<0.001), fusiform (β=0.613, p<0.001), and middle temporal (β=0.571, p<0.001) gyri (Fig. 3b).

**Figure 3.**
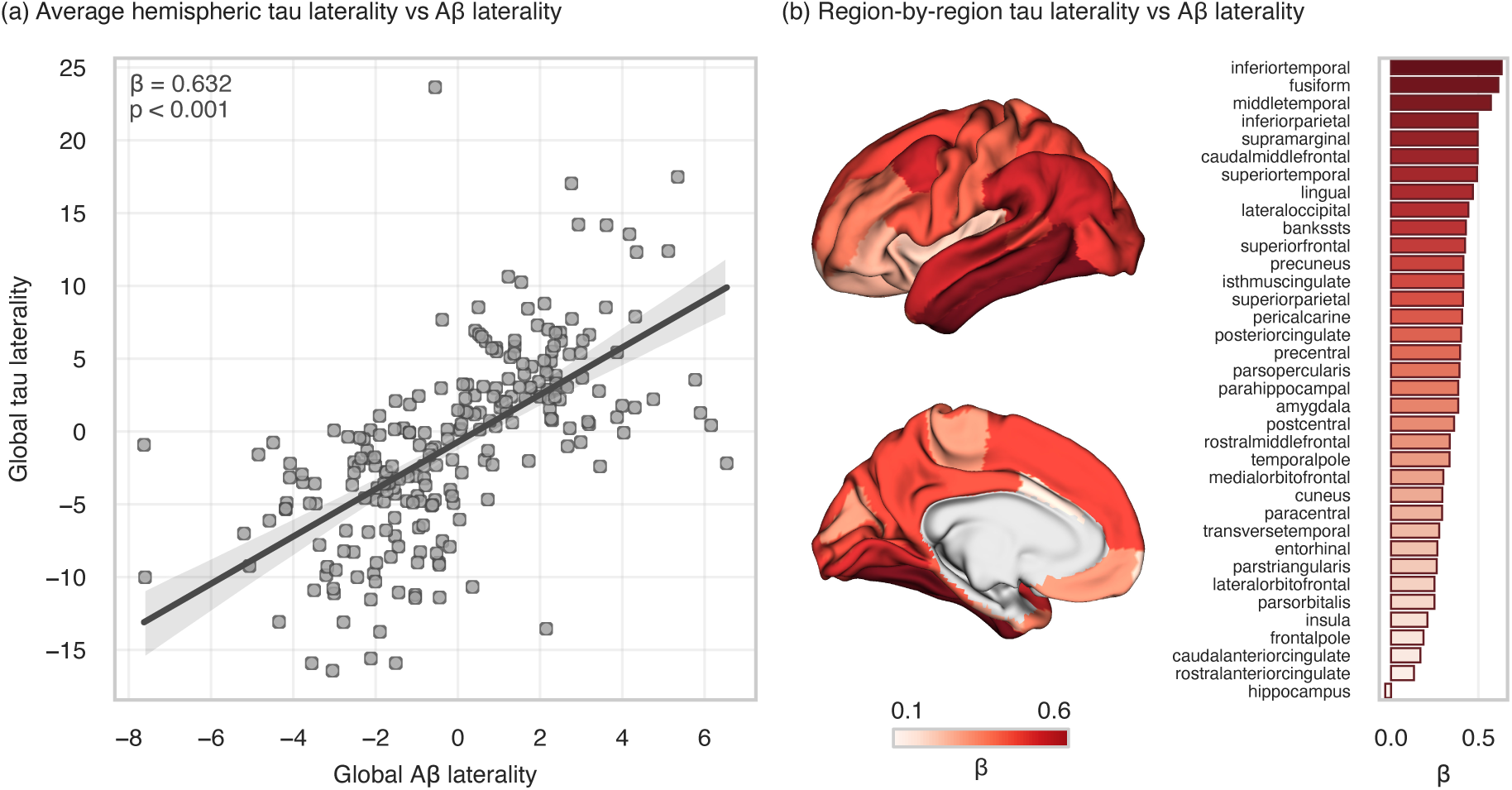
Association between asymmetry in Aβ and tau distribution: (a) Association between Aβ and tau laterality averaging the PET uptake over the whole hemisphere; (b) Regional associations between Aβ and tau laterality. Aβ – amyloid-beta

Additional analyses based on a priori selected meta-ROIs revealed the strongest associations between global Aβ laterality and tau laterality were present in a meta-ROI encompassing regions included in Braak stages III-IV (Fig S2.4; β=0.654, 95%CI=[0.555; 0.753], p<0.001). Moreover, we applied the sampled iterative local approximation algorithm (SILA) on the full BioFinder-2 cohort and found that estimated Aβ onset between the hemispheres was significantly higher within our study population’s asymmetric groups compared to the symmetric group (Fig. S2.5; see Supplementary S2 for an overview of the additional analysis performed with SILA).

### Replication in independent cohorts

To confirm the association between the laterality of Aβ and tau, we performed the same analyses on three independent cohorts – Open Access Series of Imaging Studies (OASIS-3),^29^ Anti-Amyloid Treatment in Asymptomatic Alzheimer’s Disease (A4),^30,31^ and Alzheimer’s Disease Neuroimaging Initiative (ADNI)^32^ – out of which only A+T+ subjects were included (see Supplementary Table S2.9 for demographics). We first combined the three cohorts in a single analysis controlling for the heterogeneity in the disease stages represented in the different cohorts (i.e., cognitively unimpaired or impaired). The strong relationship between asymmetries of global Aβ and tau distributions was replicated with a similar effect size compared to our main cohort (Fig. 4a; β=0.535, 95%CI=[0.425; 0.645], p<0.001). When analysed separately, each unique cohort displayed similar associations between Aβ and tau laterality despite the smaller sample sizes (Fig. 4b) – A4 (n=55; β=0.714, 95%CI=[0.516; 0.913], p<0.001), OASIS-3 (n=46; β=0.499, 95%CI=[0.208; 0.789], p=0.001), ADNI (n=133; β=0.534, 95%CI=[0.386; 0.683], p<0.001). Interestingly, all replication cohorts showed the greatest effect size of spatial relationship between the laterality of tau and Aβ pathologies in regions within Braak stages III and IV (see Supplementary Fig. S2.6).

**Figure 4.**
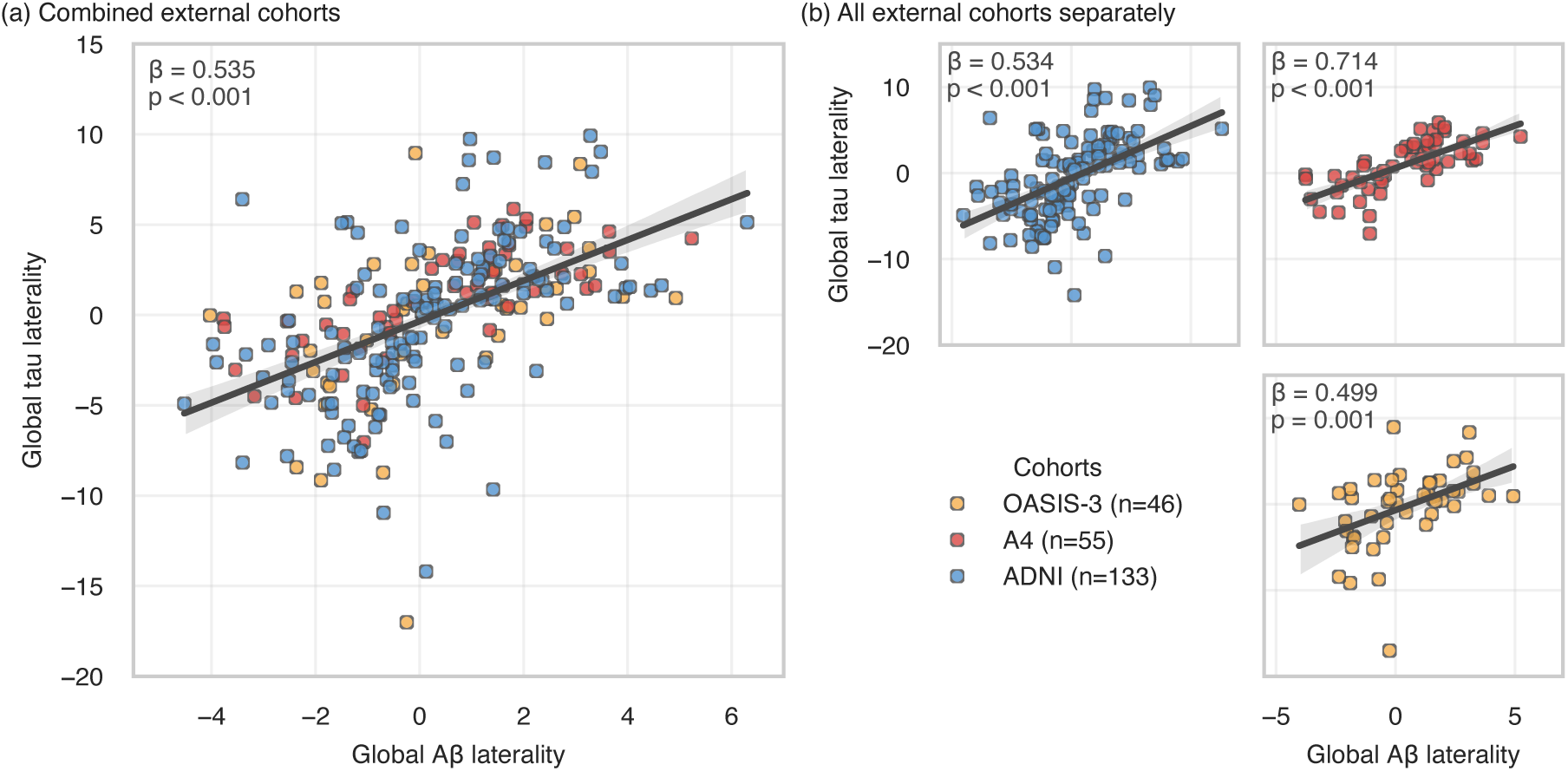
Association between global Aβ laterality and tau laterality in external cohorts: (a) All cohorts combined; (b) Cohorts separately. Aβ – amyloid-beta; OASIS-3 – Open Access Series of Imaging Studies; A4 – Anti-Amyloid Treatment in Asymptomatic Alzheimer’s Disease; ADNI – Alzheimer’s Disease Neuroimaging Initiative

### Sensitivity analyses

To support that the results did not arise from methodological issues, we performed a set of sensitivity analyses in the main cohort (see Supplementary S2 for a detailed overview). First, we investigated the association between the two pathologies using PET SUVR values corrected for partial volume effects (Fig. S2.7) and found consistent results were in all meta-ROIs (all: β>0.590, p<0.001) except Braak I-II i.e. entorhinal cortex (β=0.087, p=0.170). Second, the association between the laterality of tau and Aβ remained essentially the same when adjusting for laterality of cerebral blood flow measured with arterial spin labelling (ASL; global: β=0.498, p<0.001) or cortical thickness (global: β=0.560, p<0.001) in the model (Fig. S2.9). Moreover, Aβ-PET and tau-PET scans for nine representative cases (i.e., three cases for each asymmetrical profile) can be seen in Supplementary S3.

### Higher baseline Aβ laterality is associated with increased tau laterality over time

To further investigate the impact of baseline Aβ laterality on longitudinal change in tau laterality, we examined a subsample of 289 A+ BioFINDER-2 participants who underwent at least one Aβ-PET scan and additionally had available longitudinal tau-PET scans (range of timepoints = 2-5; average follow-up = 2.9 years). Within all these A+ subjects, a linear mixed effect (LME) model with only time as a predictor displayed an increasing tau asymmetry longitudinally (global: β=0.043, 95%CI=[0.016; 0.069], p=0.002). Next, to investigate the effects Aβ laterality on tau laterality, we included baseline Aβ laterality and covariates to the LME model. Within the same sample, higher baseline Aβ laterality was predictive of changes over time in tau in tau laterality (Fig. 5a; global: β=0.025, 95%CI=[0.003; 0.048], p=0.028), i.e., greater asymmetry of the Aβ distribution at baseline was associated with greater asymmetry of the tau distribution over time.

**Figure 5.**
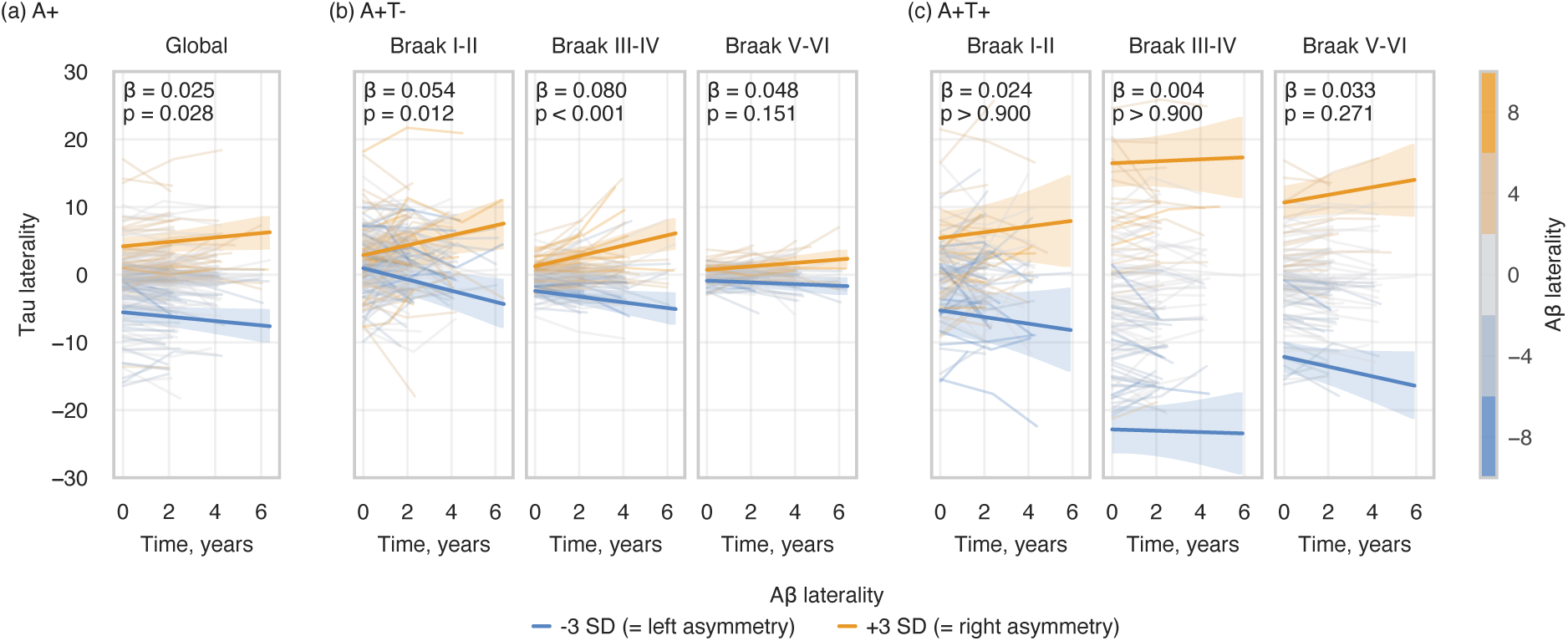
Longitudinal analysis of the association between baseline Aβ laterality and changes over time in tau laterality: (a) whole A+ sample at global meta-ROI (i.e., whole-brain for each hemisphere); (b) A+T-subsample at Braak meta-ROIs; (c) A+T+ subsample at Braak meta-ROIs. The statistical annotations indicate the effect size and significance level of the interaction between time and baseline Aβ laterality on tau laterality. For visualisation, regression lines with 95% CIs of ±3 SD baseline Aβ laterality were plotted. Aβ – amyloid-beta

We then further stratified the sample based on baseline temporal meta-ROI tau-PET uptake to A+T- (n=180) and A+T+ (n=109) groups (see Table S2.2 for demographics) and repeated the analyses according to regions defined in Braak stages. In the A+T-group (Fig. 5b), the association between Aβ laterality and tau laterality over time was confirmed in Braak stages I-II (β=0.054, 95%CI=[0.017; 0.091], p=0.012) and III-IV (β=0.080, 95%CI=[0.039; 0.120], p<0.001). In contrast, Aβ laterality was not associated with changes over time in tau laterality in the A+T+ group (Fig. 5c). For an overview of the models, see Supplementary Table S2.3, S2.4, and S2.5.

To further assess how pathological progression affects the interaction between baseline Aβ laterality and tau laterality over time, we stratified the A+T-group into additional two subgroups – individuals who remained A+T-throughout their follow-up (n=142; average follow-up = 2.9 years) and those who progressed to A+T+ during follow-up (n=38; average follow-up = 3.5 years). Subjects who maintained A+T-status (Fig. 6a) showed a significant association between baseline Aβ laterality and tau laterality over time in Braak I-II (β=0.076, 95%CI=[0.033; 0.119], p=0.002), but not in Braak III-IV or V-VI. Conversely, subjects who progressed to A+T+ (Fig. 6b) exhibited strong associations between baseline Aβ laterality and tau laterality over time in Braak III-IV (β=0.132, 95%CI=[0.077; 0.187], p<0.001) and V-VI (β=0.144, 95%CI=[0.060; 0.227], p=0.002), but not in Braak I-II. For an overview of the models, see Supplementary Table S2.6 and S2.7.

**Figure 6.**
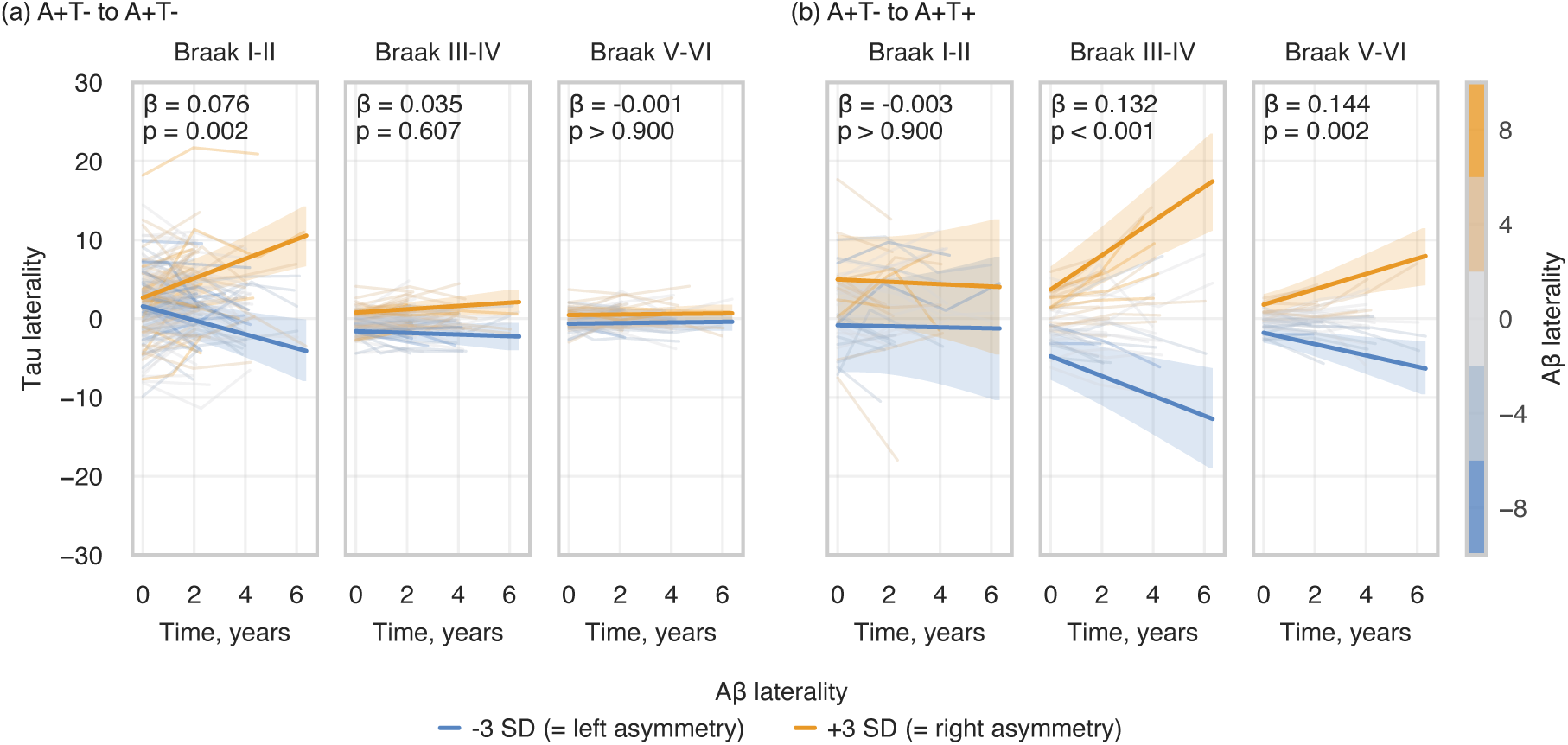
Longitudinal analysis of the association between baseline Aβ laterality and changes over time in tau laterality at Braak meta-ROIs: (a) A+T-subsample who stay A+T-throughout their follow-up; (b) A+T-subsample who progress to A+T+ during their follow-up. The statistical annotations indicate the effect size and significance level of the interaction between time and baseline Aβ laterality on tau laterality. For visualisation, regression lines with 95% CIs of ±3 SD baseline Aβ laterality were plotted. Aβ – amyloid-beta

To further confirm that these findings were not a result of our methodological choices, we repeated the analysis using partial volume corrected PET SUVR values (see Supplementary S2). Most of the findings were successfully replicated with most notably the full A+ sample exhibiting even stronger effect (Fig. S2.8; β=0.041, 95%CI=[0.021; 0.060], p<0.001) than our main analysis, but there were also some exceptions. For instance, in the sensitivity analysis the A+T+ individuals displayed a significant interaction effect in Braak V-VI (β=0.064, 95%CI=[0.034; 0.094], p<0.001), which we did not detect in the main analysis. Moreover, individuals who converted from A+T- to A+T+ did not show significant interaction of baseline Aβ laterality on tau laterality over time at Braak III-IV (β=0.064, 95%CI=[-0.012; 0.141], p=0.302) and the interaction effect was only trending towards significance in Braak V-VI after FDR-correction (β=0.093, 95%CI=[0.014; 0.171], p=0.061).

### Tau asymmetry in relation to cognitive decline

A+ participants showed distinct patterns of cognitive decline based on their tau asymmetry profiles. Higher baseline tau laterality was associated with steeper decline in modified Preclinical Alzheimer Cognitive Composite (mPACC) scores (Fig. 7a; n=259) in Braak III-IV (β=-0.134, 95%CI=[-0.172;-0.095], p<0.001) and V-VI (β=-0.157, 95%CI=[-0.194;-0.121], p<0.001). However, after adjusting for average tau uptake at the corresponding meta-ROIs, the independent effect of tau laterality on mPACC over time was statistically significant only in regions corresponding to Braak V-VI (Fig. 7b; β=-0.104, 95%CI=[-0.153;-0.055], p<0.001). In contrast, Aβ laterality did not have any significant effect on mPACC scores (Fig. 7c). See Supplementary Table S2.8 for an overview of the models.

**Figure 7.**
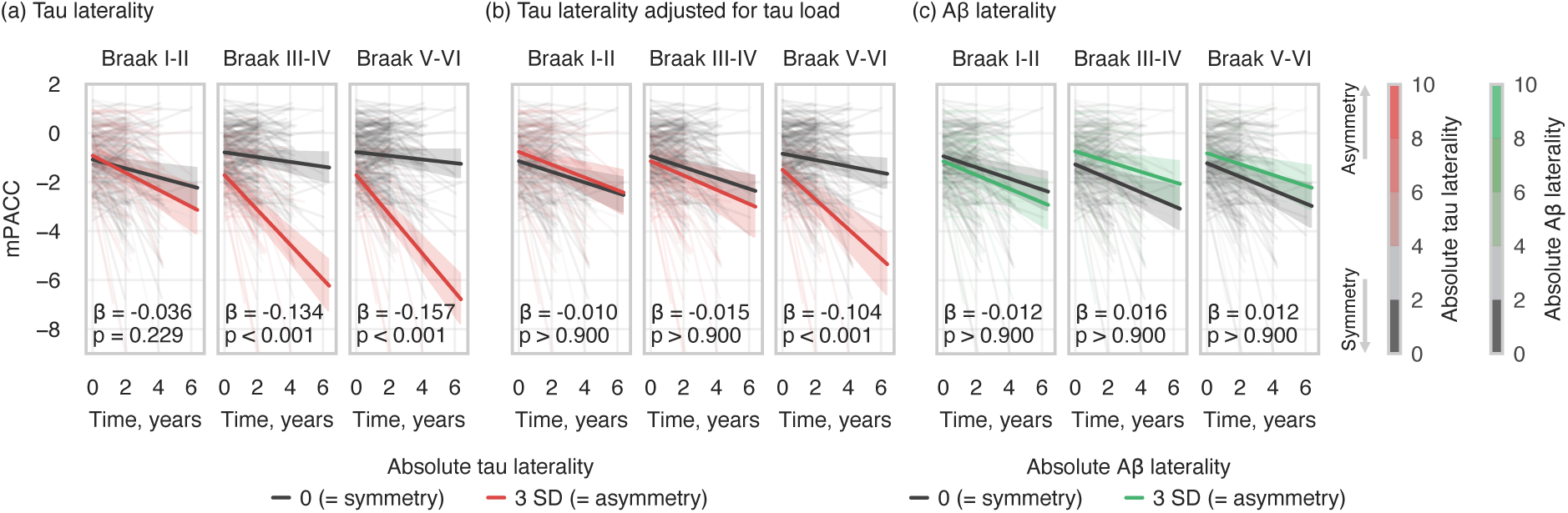
Longitudinal analysis within A+ sample over Braak meta-ROIs predicting mPACC score over time with: (a) baseline tau laterality; (b) baseline tau laterality after adjusting for tau load; (c) baseline Aβ laterality after adjusting for tau laterality, tau load, and Aβ load. Absolute values of Aβ/tau laterality were used. The statistical annotations indicate the effect size and significance level of the interaction between time and baseline Aβ/tau laterality on cognitive assessment score. For visualisation, regression lines with 95% CIs of baseline Aβ/tau laterality being 0 and 3 SD were plotted. Aβ – amyloid-beta; mPACC - modified Preclinical Alzheimer Cognitive Composite.

## Discussion

In this work, we investigated potential mechanisms underlying hemispheric asymmetry in tau distribution in AD. In contrast to one of our original hypotheses proposing reduced connectivity between hemispheres as one of the main contributing factors for asymmetrical distribution of tau, we found no evidence of an alteration in inter-hemispheric connectivity in participants with asymmetric distribution of tau compared to participants with a symmetric distribution of tau using neither functional nor structural connectivity. Moreover, the main white matter tracts connecting the two hemispheres showed similar levels of microstructural integrity assessed by fractional anisotropy and mean diffusivity. These null findings regarding inter-hemispheric connectivity might be explained by the fact that tau seeds might spread throughout the brain during the very early stages of the disease, and this process does not seem to be the rate limiting step in the build-up of tau pathology.^26^ Instead, local replication of pathogenic tau aggregates may be the main process controlling the overall rate of accumulation.^26^ This process is likely influenced by many factors including, but not limited to, Aβ pathology, which increases local soluble tau levels, and hence can accelerate the tau aggregation process ^24,25,33,34^ or possibly other mechanisms, such as impairments in brain clearance, which could affect the accumulation rate of both pathologies.^35,36^

Our results suggest that local Aβ pathology plays a critical role in determining asymmetry in tau accumulation, since tau asymmetry was preceded by asymmetry in Aβ. We systematically investigated whether this association could be explained by possible specificities in our data or analysis strategies.^37^ However, laterality patterns in tau and Aβ were strongly associated also in three replication cohorts, which included participants at different stages along the AD continuum, had varying sample sizes, and used different PET tracers. Moreover, recent investigations have confirmed the lack of affinity of tau-PET tracers to Aβ plaques, suggesting that the type of tau-PET tracer would unlikely explain the associations found.^38–40^

Another possible confounder could have been a systematic hemispheric difference in blood flow leading to an artificial difference in both Aβ and tau tracer uptake between hemispheres. However, we did not find evidence pointing in this direction. Furthermore, the association of lateralized patterns in Aβ and tau was independent from hemispheric differences in patterns of atrophy, indicating that laterality in brain atrophy did not bias the positive relationship between the two. Additional analyses using a different methodological approach (i.e., partial volume correction) further confirmed the robustness of the relationship between tau and Aβ asymmetries.

Beyond this cross-sectional relationship, the temporal dynamics of this association provide evidence for a possible mechanistic link. It is known that Aβ pathology starts to accumulate 10-30 years before cognitive impairment and precedes tau accumulation in AD.^6,24^ Most importantly, our longitudinal analyses revealed that Aβ-positive subjects with a greater degree of Aβ asymmetry at their first timepoint developed more pronounced asymmetric tau distribution over time. The relationship between Aβ and tau laterality was particularly evident in A+T-subjects, who showed strong interaction between baseline Aβ laterality and progression of tau asymmetry over time, especially in areas corresponding to early and intermediate Braak stages (I-II and III-IV). In contrast, a strong association between baseline tau and Aβ asymmetry was found in A+T+ subjects, but no association between Aβ laterality and progression over time of tau toward a more asymmetric distribution was seen. Interestingly, A+T-individuals who did not show progression in neocortical tau pathology during their follow-up exhibited a link between asymmetrical Aβ deposition and increased tau laterality over time in the entorhinal cortex (i.e., Braak I-II), whereas those whose tau distribution extended to neocortical regions during the follow-up period displayed this interaction within neocortical areas associated with later disease stages (i.e., Braak III-IV and V-VI). These results further support a role of Aβ in determining the patterns of tau distribution from the early phases of the disease course. While asymmetric distribution of Aβ pathology has been previously reported in AD,^41–45^ recent evidence from patients with comorbid epilepsy demonstrated co-lateralization of Aβ and tau in the epileptogenic hemisphere.^46^ Our findings extend this observation, suggesting that this co-occurrence of Aβ and tau is not limited to specific comorbid conditions, but represents a fundamental characteristic of AD pathophysiology.

As our findings suggest that asymmetric tau accumulation may be driven by hemispheric bias in Aβ deposits, we investigated to what extent the asymmetry in the distribution of pathology is related to cognitive decline. Consistent with previous studies investigating tau asymmetry,^7,9,12,13^ we showed that asymmetric tau distribution is associated with worse cognitive decline over time. However, this association was largely driven by tau asymmetric subjects exhibiting higher average tau load. Moreover, regardless of the association between the asymmetries of Aβ and tau, asymmetry in Aβ pathology did not have an independent effect on global cognition. These findings suggest that cognitive decline is driven by higher overall tau burden associated with asymmetrical tau distribution, rather than the asymmetrical distribution itself.

There are a few limitations to this study that should be considered. First, to provide even stronger causal roles of connectivity or Aβ deposition in determining the spatial distribution of tau pathology more data is needed from large-scale cohorts. Preferably, participants should be followed longitudinally over 10-20 years with MRI and PET scans starting many years prior to both Aβ and tau accumulation until development of substantial levels of each of the pathologies, but such cohorts are not yet available. Nevertheless, our analyses provided compelling evidence for a pivotal role of Aβ in determining the pattern of tau accumulation. Moreover, the results of the clinical trial for lecanemab, an Aβ-targeting treatment, has shown that removal of Aβ pathology leads to a reduced increase in tau PET signal over time,^47^ indicating a direct causal relationship between Aβ plaque pathology and tau accumulation. Second, as our null-finding regarding the association between inter-hemispheric connectivity and asymmetry in tau accumulation were based on macro-scale analyses, future work may benefit from assessing this relationship at finer spatial scales and employing more sophisticated approaches,^48^ which could be the key to understanding whether prion-like tau spread via connectivity drives lateralization of tau pathology. A third limitation is the relatively small number of subjects with asymmetrical tau distribution. For example, we cannot rule out that a larger sample size could have allowed us to uncover more subtle differences in structural or functional connectivity.

In summary, our study suggests that regional hemispheric vulnerability to AD pathology, especially Aβ deposits, might play a critical role in determining asymmetric distribution of tau. Specifically, asymmetric Aβ deposition appears to precede and is related to asymmetric tau accumulation, indicating that Aβ plays a critical role in the early pathophysiological cascade of AD by the suggested co-localisation of the two proteins. These results strengthen the links between Aβ and tau, supporting the hypothesis that early intervention with anti-amyloid treatments^49–51^ could help to limit the accumulation of tau pathology and downstream cognitive decline. However, further research is needed to identify the underlying mechanisms regarding the cause of one hemisphere being more susceptible to initial Aβ aggregation resulting in pathological asymmetry.

## Methods

### Main cohort

All participants within the main cohort were part of the Swedish BioFINDER-2 cohort (NCT03174938). The inclusion criteria for the present study were (1) evidence of Aβ pathology (A+); (2) age > 50 years; (3) did not fulfill the clinical criteria for other neurodegenerative diseases besides AD (e.g., frontotemporal dementia or Parkinson’s disease); (4) not diagnosed as atypical AD (e.g., posterior cortical atrophy, or primary progressive aphasia) (5) no other known severe neurological condition (e.g., brain tumour); (6) have at least one tau-PET scan. This resulted in a population of 837 participants.

In detail, cognitive assessments included mini–mental state examination (MMSE) and modified preclinical Alzheimer cognitive composite (mPACC) scores. Both clinically unimpaired and impaired individuals were included if they met the previously mentioned criteria (see Supplementary S1 for detailed inclusion and exclusion criteria). Aβ positivity was defined based on Aβ-PET neocortical uptake (SUVR > 1.033).^52^ In case Aβ-PET was not available (i.e., all the AD patients in the dementia stage, by study design), Aβ status was determined using CSF Aβ42/40 ratio measurements, with either the Roche Elecsys assay (cutoff = 0.080) or, if that was not possible, the Lumipulse G immunoassay (cutoff = 0.072).^25,53^

### Hemispheric asymmetry of pathology

For each participant, hemispheric laterality index (LI) of Aβ and tau pathologies was calculated for all brain regions and meta-ROIs using the following equation: LI (%) = 100 × (right SUVR - left SUVR) / (right SUVR + left SUVR).^10,16,54^ The cross-sectional subsample of 475 participants was selected including only Aβ positive subjects showing also evidence of tau accumulation (T+) according to tau-PET uptake in a temporal meta-ROI (i.e., covering the regions described in Braak stages I-IV) using a previously established cut-off of 1.362.^27,28^ These participants were then categorised into three groups based on the spatial distribution of tau pathology using the temporal meta-ROI tau LI – left tau asymmetric (LA) if LI < 1 SD, tau symmetric (S) if |LI| < 1 SD, or right tau asymmetric (RA) if LI > 1 SD; subjects that had LI within the ±5% range of the threshold were dropped. The resulting threshold were: |LI| > 9.70 for asymmetry and |LI| < 8.78 for symmetry. Of the 452 A+T+ subjects, 352 underwent diffusion MRI (dMRI), 318 resting-state functional MRI (RSfMRI), and 233 Aβ-PET scans. Furthermore, a second longitudinal subsample of 289 A+ participants with at least two available tau-PET scans and Aβ-PET scan at their first timepoint (i.e., baseline) were included. This subsample was stratified into A+T- (n=180) and A+T+ (n=109) based on the tau positivity at baseline (see Supplementary Table S2.2 for demographics).

All statistical analyses were performed using tableone and statsmodels packages in Python.^55,56^ Demographics were compared between the tau asymmetry groups using either one-way ANOVA, Kurskal-Wallis, or Chi-squared test depending on the type and distribution of the data. Comparisons of tau load between the groups were performed using ordinary least squares (OLS) multiple linear regressions (OLS: tau load ∼ age + sex + group). The significance level was set to p < 0.05 after Bonferroni correction accounting for the multiple comparisons between groups (i.e., p-valBonf. = p-val * 3).

### Image acquisition

#### PET

All study participants underwent PET scans on a digital GE Discovery MI scanner (General Electric Medical Systems). On this platform, tau-PET using [^18^F]RO948 (70–90 min after the injection of 365±20 MBq) and Aβ-PET using [^18^F]flutemetamol (90–110 min after the injection of ∼185 MBq) were conducted.^57^ For tau-PET, SUVR maps were calculated using the inferior cerebellar cortex as reference region.^58^ For Aβ-PET, a cortical composite SUVR was calculated using whole cerebellum as reference region.^59^ Mean SUVR values were extracted for each region of the Desikan-Killiany atlas after registering the PET images to the corresponding MRI T1-weighted scan. The average SUVR values were also calculated for meta-ROIs: global (i.e., whole brain), temporal (i.e., Braak I-IV), Braak I-II, Braak III-IV, Braak V-VI, Early-Aβ, Intermediate-Aβ, and Late-Aβ (Fig. 8; see Supplementary Table S1.1 for detailed overview).^4,5,60^

**Figure 8.**
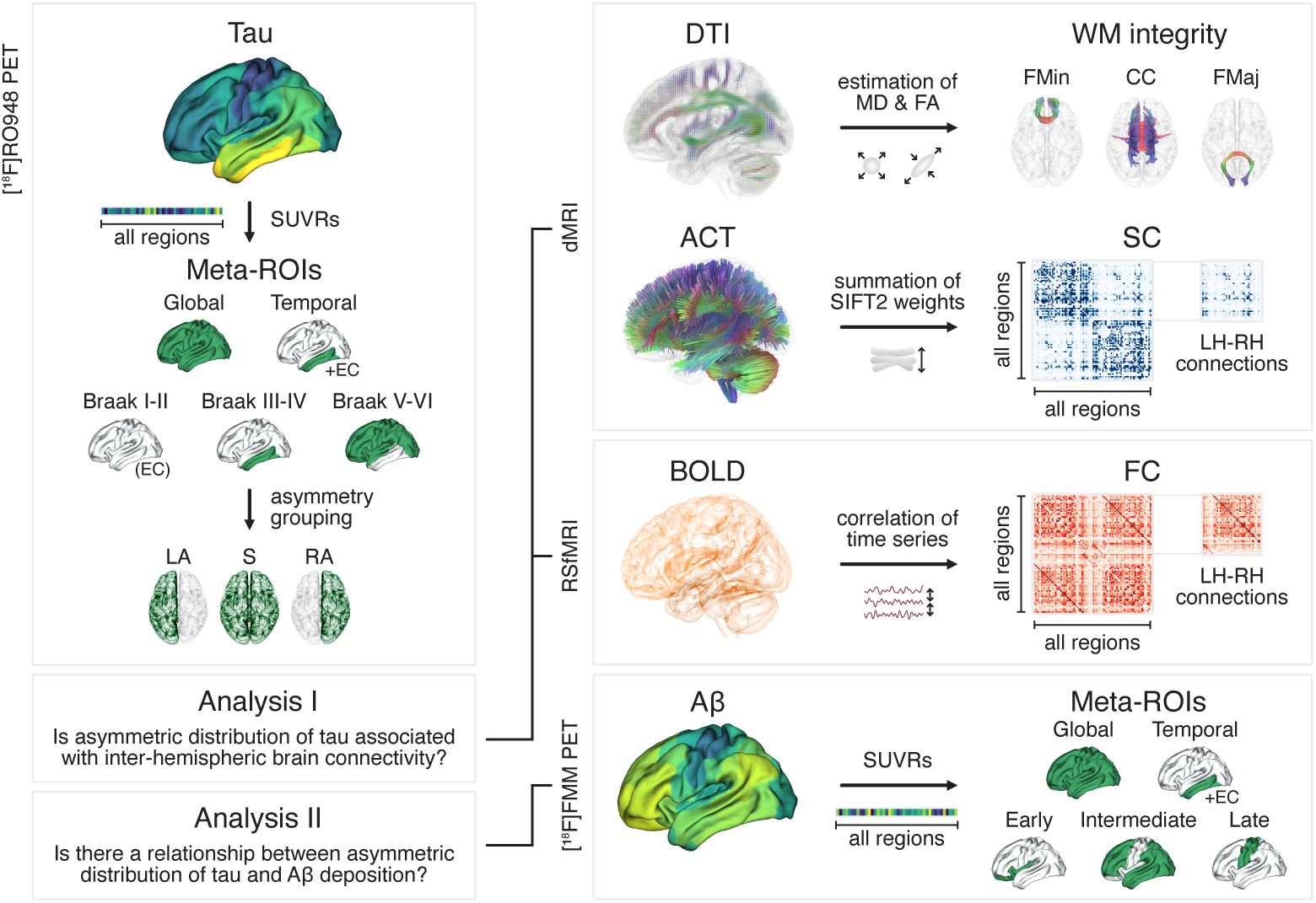
An overview of the data processing steps and analyses. PET – positron emission tomography; dMRI – diffusion magnetic resonance imaging; RSfMRI – resting-state functional magnetic resonance imaging; FMM – flutemetamol; SUVR – standardised uptake value ratio; Meta-ROI – meta region of interest; EC – entorhinal cortex; LA – left tau asymmetric; S – tau symmetric; RA – right tau asymmetric; DTI – diffusion tensor imaging; MD – mean diffusivity; FA – fractional anisotropy; FMin – Forceps Minor; CC – Corpus Callosum; FMaj – Forceps Major; ACT – anatomically constrained tractography; SIFT2 – spherical-deconvolution informed filtering of tractograms; LH/RH – left/right hemisphere; SC – structural connectivity; BOLD – blood-oxygen-level-dependent imaging; FC – functional connectivity; Aβ – amyloid-beta

#### MRI protocol

The MRI imaging was conducted using a MAGNETOM Prisma 3T MRI scanner (Siemens Healthineers) with a 64-channel head coil. RSfMRI was acquired using a gradient-echo planar sequence (eyes closed; in-plane resolution = 3×3mm^2^; slice thickness = 3.6mm; repetition time = 1020ms; echo time = 30ms; flip-angle = 63°; 462 dynamic scans over a period of 7.85min). For dMRI, 104 diffusion-weighted imaging volumes were acquired using a single-shot echo-planar imaging sequence (repetition time = 3500ms; echo time = 73ms; resolution = 2×2×2mm^3^; field of view = 220×220×124mm^3^; b-values range = 0, 100, 1000 and 2500s/mm^2^ distributed over 2, 6, 32 and 64 directions; 2-fold parallel acceleration and partial Fourier factor = 7/8). T1-weighted structural images were also acquired using a magnetization-prepared rapid gradient-echo (MPRAGE) sequence (inversion time = 1100ms; flip-angle = 9°; echo time = 2.54ms; echo spacing = 7.3ms; repetition time = 1900ms; receiver bandwidth = 220 Hz/pixel; voxel size = 1×1×1mm^3^). Generalized autocalibrating partially parallel acquisitions (GRAPPA) was applied with an acceleration factor of 2 and 24 reference lines. Additionally, ASL scans were acquired on a subset of the sample using a prototype 3D pseudo-continuous (pCASL) sequence with background suppression and gradient- and spin-echo (GRASE) readout as described previously.^61^

#### MRI pre-processing

T1-weighted structural MPRAGE images were pre-processed using FreeSurfer (version 6.0, https://surfer.nmr.mgh.harvard.edu).^62^ This included various steps, such as correction for intensity homogeneity, skull stripping, and tissue segmentation. Diffusion MRI images were preprocessed using a combination of FSL (FMRIB Software Library, version 6.0.4; Oxford, UK) and MRtrix3 tools.^63,64^ These images underwent correction for susceptibility-induced distortions, using images acquired with opposite phase polarities, motion, and Eddy current induced artifacts. RSfMRI data underwent preprocessing using Configurable Pipeline for the Analysis of Connectomes (C-PAC), including motion correction, bandpass filtering (0.01-0.1 Hz), and noise regression.^65^ Susceptibility distortion was corrected using T2-based unwarping, and outlier frames identified by DVARS (D refers to temporal derivative of the time courses and VARS refers to RMS variance over voxels) were censored.

#### Estimation of MRI-based connectivity measures

Structural connectivity (SC) and functional connectivity (FC) were estimated using a combination of MRtrix3, FSL, FreeSurfer, and Nilearn software packages, integrated through NiPype interfaces in Python.^62–64,66,67^ See Supplementary S1 for a more detailed description of the process.

In brief, FC matrices were derived from pre-processed subject-space RSfMRI data using Pearson correlation after Fisher’s z-transformation,^68^ and estimated for 84 cortical and sub-cortical regions defined in the Desikan-Killiany atlas and in the ASEG protocol.^69,70^ For SC, response functions for white matter (WM), grey matter (GM), and cerebrospinal fluid (CSF) were estimated using the "dhollander" algorithm^71^ on pre-processed diffusion MRI data from 60 cognitively unimpaired A-T- and 40 A+T-participants of the BioFINDER-2 cohort, and fiber orientation distributions (FOD) were derived via multi-shell multi-tissue constrained spherical deconvolution.^72^ Anatomically-constrained tractography (ACT) with the "iFOD2" algorithm generated ten million streamlines, which were filtered using SIFT2 to reduce overestimation bias.^73–76^ SC matrices were constructed for the same 84 regions employed for FC,^69^ with summation of SIFT2-weighted streamlines per region used as edges.

For evaluating microstructural integrity, the diffusion tensor imaging (DTI) model was fitted to the diffusion MRI data using the volumes acquired with a b-value up to 1000 s/mm², and fractional anisotropy (FA) and mean diffusivity (MD) maps were derived^77,78^. The main inter-hemispheric white matter tracts – corpus callosum, forceps major, forceps minor – were segmented using TractSeg and the mean FA and MD values were extracted for each tract.^79^

### Comparison of brain connectivity between tau asymmetry groups

#### Inter-hemispheric connectivity and microstructural integrity

To reduce noise and focus on the most biologically plausible connections, we masked both the FC and SC matrices for all subjects by selecting only the top 10% inter-hemispheric connections identified in 294 age-matched healthy controls with no evidence of Aβ or tau pathology from the BioFINDER-2 cohort.

For each subject, we then calculated the global inter-hemispheric FC and SC by averaging the connections between all the left nodes connecting to the right (Fig. 8). These averages were subsequently compared between the tau asymmetry groups (i.e., LA vs S, RA vs S, and LA vs RA) in the cross-sectional A+T+ sample using OLS multiple linear regressions (OLS: FC/SC ~ age + sex + average global tau load + group) with the significance level set to p < 0.05 after Bonferroni correction accounting for the multiple comparisons between groups (i.e., p-valBonf. = p-val * 3). Additionally, FC and SC between each counter-lateral regions were compared between the groups with the same statistical model but with the significance level set to p < 0.05 after applying False Discovery Rate (FDR) correction (Benjamini & Hochberg method). Average FA and MD values in the main tracts connecting the two hemispheres were also compared between the groups, similarly to FC and SC.

#### Whole-brain connectomics

Network Based Statistic (NBS) method^80^ was applied on FC and SC matrices to compare the whole-brain connectome differences between the tau asymmetry groups. In short, NBS is a cluster-based method that consists of three main steps: (1) edges that, when compared between groups, surpass a given statistical threshold (e.g., t=3.0) are identified, (2) components (i.e., connected subgraphs or clusters of topologically contiguous supra-threshold edges) are detected, and (3) permutation testing adjusting for family-wise error (FWE) is performed to assign a p-value for each detected component based on its size relative to the null distribution of component sizes obtained through permutation. It is often recommended to repeat NBS using different statistical thresholds in step (1).^80^ This study used thresholds of 2.5, 3.0, and 3.5. Moreover, the comparisons were adjusted for age, sex, and average global tau load. 5000 permutations were performed, and the significance level was set to p < 0.05 after FWE correction.

### Association between the distribution of Aβ and tau pathologies

The association between the laterality of Aβ and tau (i.e., Aβ LI vs tau LI) was investigated in the cross-sectional A+T+ sample for each region defined in the Desikan-Killiany atlas and each meta-ROI described above (Fig. 8). The statistical analyses were performed using OLS multiple linear regressions (OLS: tau LI ∼ age + sex + Aβ LI) with the significance level set to p < 0.05 for the meta-ROI based analyses and FDR-corrected for region-by-region analyses.

The association between baseline Aβ laterality and tau laterality over time was investigated in the longitudinal A+, A+T-, and A+T+ samples for global, Braak I-II, Braak III-IV, and Braak V-VI meta-ROIs. The statistical analyses were performed using Linear Mixed Effects (LME) models with random intercepts and slopes for time and participants (LME: tau LI ∼ time * (agebaseline + sex + Aβ LIbaseline) + [1 + time | participant]), with the significance level set to p < 0.05 after Bonferroni correction for the number of comparisons performed (i.e., number of meta-ROIs).

### Association between the distribution of pathologies and cognition

The association between baseline Aβ and tau laterality and cognitive decline was investigated in the longitudinal A+ sample for Braak I-II, Braak III-IV, and Braak V-VI meta-ROIs. The statistical analyses were performed similarly using LME modelling with random intercepts and slopes for time and participants, but with three different models: (1) a base model testing tau laterality effects on cognition (LME: mPACC ∼ time * (agebaseline + sex + tau LIbaseline) + [1 + time | participant]), (2) a model testing tau laterality effects on cognition but controlling for tau load (LME: mPACC ∼ time * (agebaseline + sex + tau loadbaseline + tau LIbaseline) + [1 + time | participant]), and (3) a model testing Aβ laterality effects on cognition (LME: mPACC time * (agebaseline + sex + Aβ LIbaseline) + [1 + time | participant]). The significance level for all models was set to p < 0.05 after Bonferroni correction for the number of comparisons performed (i.e., number of meta-ROIs).

### Replication in independent cohorts

Three external cohorts (see Table S2.9 for demographics) were used to validate the association between the laterality of Aβ and tau distribution – Open Access Series of Imaging Studies (OASIS-3; https://sites.wustl.edu/oasisbrains/), Anti-Amyloid Treatment in Asymptomatic Alzheimer’s Disease (A4; https://www.a4studydata.org/), and Alzheimer’s Disease Neuroimaging Initiative (ADNI; https://adni.loni.usc.edu/). For each cohort, only subjects with evidence of both Aβ and tau (A+T+) and with at least one Aβ-PET and one tau-PET scan available were selected. All these cohorts used [18F]florbetapir PET for Aβ imaging and [18F]flortaucipir PET for tau imaging. Aβ positivity was pre-defined by the cohort creators for OASIS-3 and ADNI datasets, but a cut-off of Global Aβ-PET SUVR > 1.11 was used for A4.^81^ Tau positivity was defined based on a Temporal meta-ROI SUVR > 1.34 cut-off for each external cohort.^82^

## Data availability

Four different cohorts were used in this study: BioFINDER-2, ADNI, A4, and OASIS-3. For BioFINDER-2 data, anonymized data will be shared by request from a qualified academic investigator as long as data transfer is in agreement with European Union legislation on the general data protection regulation and decisions by the Swedish Ethical Review Authority and Region Skåne, which should be regulated in a material transfer agreement. ADNI, A4 and OASIS3 are publicly available datasets and can be obtained from http://adni.loni.usc.edu/, https://ida.loni.usc.edu/ and https://sites.wustl.edu/oasisbrains/, respectively.

## Code availability

All the necessary code for the analyses conducted within this study is available on GitHub (https://github.com/teanijarv/phd-neurodegeneration-imaging/tree/main/projects/01_tau_asymmetry).

## Acknowledgements

We would like to express our gratitude to the research volunteers who participated in the studies from which these data were obtained and their supportive families.

## Author information

### Authors and Affiliations

**Clinical Memory Research Unit, Department of Clinical Sciences Malmö, Faculty of Medicine, Lund University, Lund, Sweden**

Toomas Erik Anijärv, Rik Ossenkoppele, Ruben Smith, Alexa Pichet Binette, Lyduine E. Collij, Harry H. Behjat, Linda Karlsson, Olof Strandberg, Erik Stomrud, Sebastian Palmqvist, Niklas Mattsson-Carlgren, Nicola Spotorno & Oskar Hansson

**Alzheimer Center Amsterdam, Neurology, Vrije Universiteit Amsterdam, Amsterdam UMC, Amsterdam, the Netherlands**

Rik Ossenkoppele

**Neurodegeneration, Amsterdam Neuroscience, Amsterdam, the Netherlands**

Rik Ossenkoppele

**Memory Clinic, Skåne University Hospital, Malmö, Sweden**

Ruben Smith, Erik Stomrud, Sebastian Palmqvist, Niklas Mattsson-Carlgren

**Department of Physiology and Pharmacology, Université de Montréal, Montréal, Quebec, Canada**

Alexa Pichet Binette

**Centre de Recherche de l’Institut Universitaire de Gériatrie de Montréal, Montréal, Quebec, Canada**

Alexa Pichet Binette

**Radiology and Nuclear Medicine, Vrije Universiteit Amsterdam, Amsterdam UMC, Amsterdam, the Netherlands**

Lyduine E. Collij

**Brain Imaging, Amsterdam Neuroscience, Amsterdam, the Netherlands**

Lyduine E. Collij

**SciLifeLab, Department of Clinical Sciences Malmö, Faculty of Medicine, Lund University, Lund, Sweden**

Jonathan Rittmo & Jacob W. Vogel

**Department of Neuropsychology, Ruhr University Bochum, Bochum, Germany**

Khazar Ahmadi

**Diagnostic Radiology, Institution for Clinical Sciences, Lund University, Lund, Sweden**

Danielle van Westen

**Wallenberg Center for Molecular Medicine, Lund University, Lund, Sweden**

Niklas Mattsson-Carlgren

### Consortia

#### Alzheimer’s Disease Neuroimaging Initiative

Michael Weiner, Paul Aisen, Ronald Petersen, Clifford R. Jack Jr., William Jagust, John Q. Trojanowki, Arthur W. Toga, Laurel Beckett, Robert C. Green, Andrew J. Saykin, John Morris, Leslie M. Shaw, Enchi Liu, Tom Montine, Ronald G. Thomas, Michael Donohue, Sarah Walter, Devon Gessert, Tamie Sather, Gus Jiminez, Danielle Harvey, Michael Donohue, Matthew Bernstein, Nick Fox, Paul Thompson, Norbert Schuff, Charles DeCArli, Bret Borowski, Jeff Gunter, Matt Senjem, Prashanthi Vemuri, David Jones, Kejal Kantarci, Chad Ward, Robert A. Koeppe, Norm Foster, Eric M. Reiman, Kewei Chen, Chet Mathis, Susan Landau, Nigel J. Cairns, Erin Householder, Lisa Taylor Reinwald, Virginia Lee, Magdalena Korecka, Michal Figurski, Karen Crawford, Scott Neu, Tatiana M. Foroud, Steven Potkin, Li Shen, Faber Kelley, Sungeun Kim, Kwangsik Nho, Zaven Kachaturian, Richard Frank, Peter J. Snyder, Susan Molchan, Jeffrey Kaye, Joseph Quinn, Betty Lind, Raina Carter, Sara Dolen, Lon S. Schneider, Sonia Pawluczyk, Mauricio Beccera, Liberty Teodoro, Bryan M. Spann, James Brewer, Helen Vanderswag, Adam Fleisher, Judith L. Heidebrink, Joanne L. Lord, Ronald Petersen, Sara S. Mason, Colleen S. Albers, David Knopman, Kris Johnson, Rachelle S. Doody, Javier Villanueva Meyer, Munir Chowdhury, Susan Rountree, Mimi Dang, Yaakov Stern, Lawrence S. Honig, Karen L. Bell, Beau Ances, John C. Morris, Maria Carroll, Sue Leon, Erin Householder, Mark A. Mintun, Stacy Schneider, Angela OliverNG, Randall Griffith, David Clark, David Geldmacher, John Brockington, Erik Roberson, Hillel Grossman, Effie Mitsis, Leyla deToledo-Morrell, Raj C. Shah, Ranjan Duara, Daniel Varon, Maria T. Greig, Peggy Roberts, Marilyn Albert, Chiadi Onyike, Daniel D’Agostino II, Stephanie Kielb, James E. Galvin, Dana M. Pogorelec, Brittany Cerbone, Christina A. Michel, Henry Rusinek, Mony J. de Leon, Lidia Glodzik, Susan De Santi, P. Murali Doraiswamy, Jeffrey R. Petrella, Terence Z. Wong, Steven E. Arnold, Jason H. Karlawish, David Wolk, Charles D. Smith, Greg Jicha, Peter Hardy, Partha Sinha, Elizabeth Oates, Gary Conrad, Oscar L. Lopez, MaryAnn Oakley, Donna M. Simpson, Anton P. Porsteinsson, Bonnie S. Goldstein, Kim Martin, Kelly M. Makino, M. Saleem Ismail, Connie Brand, Ruth A. Mulnard, Gaby Thai, Catherine Mc Adams Ortiz, Kyle Womack, Dana Mathews, Mary Quiceno, Ramon Diaz Arrastia, Richard King, Myron Weiner, Kristen Martin Cook, Michael DeVous, Allan I. Levey, James J. Lah, Janet S. Cellar, Jeffrey M. Burns, Heather S. Anderson, Russell H. Swerdlow, Liana Apostolova, Kathleen Tingus, Ellen Woo, Daniel H. S. Silverman, Po H. Lu, George Bartzokis, Neill R. Graff Radford, Francine ParfittH, Tracy Kendall, Heather Johnson, Martin R. Farlow, Ann Marie Hake, Brandy R. Matthews, Scott Herring, Cynthia Hunt, Christopher H. van Dyck, Richard E. Carson, Martha G. MacAvoy, Howard Chertkow, Howard Bergman, Chris Hosein, Sandra Black, Bojana Stefanovic, Curtis Caldwell, Ging Yuek Robin Hsiung, Howard Feldman, Benita Mudge, Michele Assaly Past, Andrew Kertesz, John Rogers, Dick Trost, Charles Bernick, Donna Munic, Diana Kerwin, Marek Marsel Mesulam, Kristine Lipowski, Chuang Kuo Wu, Nancy Johnson, Carl Sadowsky, Walter Martinez, Teresa Villena, Raymond Scott Turner, Kathleen Johnson, Brigid Reynolds, Reisa A. Sperling, Keith A. Johnson, Gad Marshall, Meghan Frey, Jerome Yesavage, Joy L. Taylor, Barton Lane, Allyson Rosen, Jared Tinklenberg, Marwan N. Sabbagh, Christine M. Belden, Sandra A. Jacobson, Sherye A. Sirrel, Neil Kowall, Ronald Killiany, Andrew E. Budson, Alexander Norbash, Patricia Lynn Johnson, Thomas O. Obisesan, Saba Wolday, Joanne Allard, Alan Lerner, Paula Ogrocki, Leon Hudson, Evan Fletcher, Owen Carmichael, John Olichney, Charles DeCarli, Smita Kittur, Michael Borrie, T. Y. Lee, Rob Bartha, Sterling Johnson, Sanjay Asthana, Cynthia M. Carlsson, Steven G. Potkin, Adrian Preda, Dana Nguyen, Pierre Tariot, Adam Fleisher, Stephanie Reeder, Vernice Bates, Horacio Capote, Michelle Rainka, Douglas W. Scharre, Maria Kataki, Anahita Adeli, Earl A. Zimmerman, Dzintra Celmins, Alice D. Brown, Godfrey D. Pearlson, Karen Blank, Karen Anderson, Robert B. Santulli, Tamar J. Kitzmiller, Eben S. Schwartz, Kaycee M. SinkS, Jeff D. Williamson, Pradeep Garg, Franklin Watkins, Brian R. Ott, Henry Querfurth, Geoffrey Tremont, Stephen Salloway, Paul Malloy, Stephen Correia, Howard J. Rosen, Bruce L. Miller, Jacobo Mintzer, Kenneth Spicer, David Bachman, Elizabether Finger, Stephen Pasternak, Irina Rachinsky, John Rogers, Andrew Kertesz, Dick Drost, Nunzio Pomara, Raymundo Hernando, Antero Sarrael, Susan K. Schultz, Laura L. Boles Ponto, Hyungsub Shim, Karen Elizabeth Smith, Norman Relkin, Gloria Chaing, Lisa Raudin, Amanda Smith, Kristin Fargher & Balebail Ashok Raj

### Contributions

T.E.A., N.S., and O.H. designed the study. T.E.A. and N.S. had full access to raw data. T.E.A. performed data processing and carried out the statistical analyses. T.E.A. wrote the manuscript and had the final responsibility to submit for publication. N.S. and O.H. contributed as the main supervisors of the work. All other authors contributed demographic, clinical, biomarker and neuroimaging data, contributed to the interpretation of the results and critically reviewed the manuscript.

### Corresponding authors

Correspondence to Toomas Erik Anijärv, Nicola Spotorno, or Oskar Hansson.

## References

1. Ferreira D, Nordberg A, Westman E. Biological subtypes of Alzheimer disease. Neurology. 2020;94(10):436–448. doi:10.1212/WNL.0000000000009058

2. Baumeister H, Vogel JW, Insel PS, et al. A generalizable data-driven model of atrophy heterogeneity and progression in memory clinic settings. Brain. 2024;147(7):2400–2413. doi:10.1093/brain/awae118

3. Šimić G, Babić Leko M, Wray S, et al. Tau Protein Hyperphosphorylation and Aggregation in Alzheimer’s Disease and Other Tauopathies, and Possible Neuroprotective Strategies. Biomolecules. 2016;6(1):6. doi:10.3390/biom6010006

4. Braak H, Braak E. Neuropathological stageing of Alzheimer-related changes. Acta Neuropathol (Berl). 1991;82(4):239–259. doi:10.1007/BF00308809

5. Braak H, Alafuzoff I, Arzberger T, Kretzschmar H, Del Tredici K. Staging of Alzheimer disease-associated neurofibrillary pathology using paraffin sections and immunocytochemistry. Acta Neuropathol (Berl). 2006;112(4):389–404. doi:10.1007/s00401-006-0127-z

6. Hansson O. Biomarkers for neurodegenerative diseases. Nat Med. 2021;27(6):954–963. doi:10.1038/s41591-021-01382-x

7. Vogel JW, Young AL, Oxtoby NP, et al. Four distinct trajectories of tau deposition identified in Alzheimer’s disease. Nat Med. 2021;27(5):871–881. doi:10.1038/s41591-021-01309-6

8. Ossenkoppele R, Kant R van der, Hansson O. Tau biomarkers in Alzheimer’s disease: towards implementation in clinical practice and trials. Lancet Neurol. 2022;21(8):726–734. doi:10.1016/S1474-4422(22)00168-5

9. Lu J, Zhang Z, Wu P, et al. The heterogeneity of asymmetric tau distribution is associated with an early age at onset and poor prognosis in Alzheimer’s disease. NeuroImage Clin. 2023;38:103416. doi:10.1016/j.nicl.2023.103416

10. Young CB, Winer JR, Younes K, et al. Divergent Cortical Tau Positron Emission Tomography Patterns Among Patients With Preclinical Alzheimer Disease. JAMA Neurol. 2022;79(6):592–603. doi:10.1001/jamaneurol.2022.0676

11. Younes K, Smith V, Johns E, et al. Temporal tau asymmetry spectrum influences divergent behavior and language patterns in Alzheimer‘s disease. Brain Behav Immun. 2024;119:807–817. doi:10.1016/j.bbi.2024.05.002

12. King A, Bodi I, Nolan M, Troakes C, Al-Sarraj S. Assessment of the degree of asymmetry of pathological features in neurodegenerative diseases. What is the significance for brain banks? J Neural Transm. 2015;122(10):1499–1508. doi:10.1007/s00702-015-1410-8

13. Tremblay C, Serrano GE, Intorcia AJ, et al. Hemispheric Asymmetry and Atypical Lobar Progression of Alzheimer-Type Tauopathy. J Neuropathol Exp Neurol. 2022;81(3):158–171. doi:10.1093/jnen/nlac008

14. Crutch SJ, Schott JM, Rabinovici GD, et al. Consensus classification of posterior cortical atrophy. Alzheimers Dement. 2017;13(8):870–884. doi:10.1016/j.jalz.2017.01.014

15. Gorno-Tempini ML, Hillis AE, Weintraub S, et al. Classification of primary progressive aphasia and its variants. Neurology. 2011;76(11):1006–1014. doi:10.1212/WNL.0b013e31821103e6

16. Ossenkoppele R, Schonhaut DR, Schöll M, et al. Tau PET patterns mirror clinical and neuroanatomical variability in Alzheimer’s disease. Brain. 2016;139(5):1551–1567. doi:10.1093/brain/aww027

17. Tetzloff KA, Graff-Radford J, Martin PR, et al. Regional Distribution, Asymmetry, and Clinical Correlates of Tau Uptake on [ 18F]AV-1451 PET in Atypical Alzheimer’s Disease. J Alzheimers Dis. 2018;62(4):1713–1724. doi:10.3233/JAD-170740

18. Martersteck A, Ayala I, Ohm DT, et al. Focal amyloid and asymmetric tau in an imaging-to-autopsy case of clinical primary progressive aphasia with Alzheimer disease neuropathology. Acta Neuropathol Commun. 2022;10(1):111. doi:10.1186/s40478-022-01412-w

19. Vogel JW, Iturria-Medina Y, Strandberg OT, et al. Spread of pathological tau proteins through communicating neurons in human Alzheimer’s disease. Nat Commun. 2020;11(1):2612. doi:10.1038/s41467-020-15701-2

20. Yang F, Chowdhury SR, Jacobs HIL, et al. Longitudinal predictive modeling of tau progression along the structural connectome. NeuroImage. 2021;237:118126. doi:10.1016/j.neuroimage.2021.118126

21. Franzmeier N, Neitzel J, Rubinski A, et al. Functional brain architecture is associated with the rate of tau accumulation in Alzheimer’s disease. Nat Commun. 2020;11(1):347. doi:10.1038/s41467-019-14159-1

22. Steward A, Biel D, Brendel M, et al. Functional network segregation is associated with attenuated tau spreading in Alzheimer’s disease. Alzheimers Dement. 2023;19(5):2034–2046. doi:10.1002/alz.12867

23. Cope TE, Rittman T, Borchert RJ, et al. Tau burden and the functional connectome in Alzheimer’s disease and progressive supranuclear palsy. Brain. 2018;141(2):550–567. doi:10.1093/brain/awx347

24. Mattsson-Carlgren N, Andersson E, Janelidze S, et al. Aβ deposition is associated with increases in soluble and phosphorylated tau that precede a positive Tau PET in Alzheimer’s disease. Sci Adv. 2020;6(16):eaaz2387. doi:10.1126/sciadv.aaz2387

25. Pichet Binette A, Franzmeier N, Spotorno N, et al. Amyloid-associated increases in soluble tau relate to tau aggregation rates and cognitive decline in early Alzheimer’s disease. Nat Commun. 2022;13:6635. doi:10.1038/s41467-022-34129-4

26. Meisl G, Hidari E, Allinson K, et al. In vivo rate-determining steps of tau seed accumulation in Alzheimer’s disease. Sci Adv. 2021;7(44):eabh1448. doi:10.1126/sciadv.abh1448

27. Cho H, Choi JY, Hwang MS, et al. In vivo cortical spreading pattern of tau and amyloid in the Alzheimer disease spectrum. Ann Neurol. 2016;80(2):247–258. doi:10.1002/ana.24711

28. Leuzy A, Smith R, Ossenkoppele R, et al. Diagnostic Performance of RO948 F 18 Tau Positron Emission Tomography in the Differentiation of Alzheimer Disease From Other Neurodegenerative Disorders. JAMA Neurol. 2020;77(8):955–965. doi:10.1001/jamaneurol.2020.0989

29. LaMontagne PJ, Benzinger TL, Morris JC, et al. OASIS-3: Longitudinal Neuroimaging, Clinical, and Cognitive Dataset for Normal Aging and Alzheimer Disease. Published online December 15, 2019:2019.12.13.19014902. doi:10.1101/2019.12.13.19014902

30. Sperling RA, Rentz DM, Johnson KA, et al. The A4 Study: Stopping AD Before Symptoms Begin? Sci Transl Med. 2014;6(228):228fs13-228fs13. doi:10.1126/scitranslmed.3007941

31. Insel PS, Donohue MC, Sperling R, Hansson O, Mattsson-Carlgren N. The A4 study: β-amyloid and cognition in 4432 cognitively unimpaired adults. Ann Clin Transl Neurol. 2020;7(5):776–785. doi:10.1002/acn3.51048

32. Petersen RC, Aisen PS, Beckett LA, et al. Alzheimer’s Disease Neuroimaging Initiative (ADNI). Neurology. 2010;74(3):201–209. doi:10.1212/WNL.0b013e3181cb3e25

33. Bennett RE, DeVos SL, Dujardin S, et al. Enhanced Tau Aggregation in the Presence of Amyloid β. Am J Pathol. 2017;187(7):1601–1612. doi:10.1016/j.ajpath.2017.03.011

34. Kwak SS, Washicosky KJ, Brand E, et al. Amyloid-β42/40 ratio drives tau pathology in 3D human neural cell culture models of Alzheimer’s disease. Nat Commun. 2020;11(1):1377. doi:10.1038/s41467-020-15120-3

35. Tarasoff-Conway JM, Carare RO, Osorio RS, et al. Clearance systems in the brain— implications for Alzheimer disease. Nat Rev Neurol. 2015;11(8):457–470. doi:10.1038/nrneurol.2015.119

36. Ishida K, Yamada K, Nishiyama R, et al. Glymphatic system clears extracellular tau and protects from tau aggregation and neurodegeneration. J Exp Med. 2022;219(3):e20211275. doi:10.1084/jem.20211275

37. Lemoine L, Leuzy A, Chiotis K, Rodriguez-Vieitez E, Nordberg A. Tau positron emission tomography imaging in tauopathies: The added hurdle of off-target binding. Alzheimers Dement Diagn Assess Dis Monit. 2018;10:232. doi:10.1016/j.dadm.2018.01.007

38. Kuwabara H, Comley RA, Borroni E, et al. Evaluation of 18F-RO-948 PET for Quantitative Assessment of Tau Accumulation in the Human Brain. J Nucl Med. 2018;59(12):1877. doi:10.2967/jnumed.118.214437

39. Freiburghaus T, Pawlik D, Oliveira Hauer K, et al. Association of in vivo retention of [18f] flortaucipir pet with tau neuropathology in corresponding brain regions. Acta Neuropathol (Berl). 2024;148(1):44. doi:10.1007/s00401-024-02801-2

40. Smith R, Wibom M, Pawlik D, Englund E, Hansson O. Correlation of In Vivo [18F]Flortaucipir With Postmortem Alzheimer Disease Tau Pathology. JAMA Neurol. 2019;76(3):310–317. doi:10.1001/jamaneurol.2018.3692

41. Yoon HJ, Kim BS, Jeong JH, Kim GH, Park HK, Chun MY. Asymmetric Amyloid Deposition as an Early Sign of Progression in Mild Cognitive Impairment Due to Alzheimer Disease. Clin Nucl Med. 2021;46(7):527–531. doi:10.1097/RLU.0000000000003662

42. Kjeldsen PL, Parbo P, Hansen KV, et al. Asymmetric amyloid deposition in preclinical Alzheimer’s disease: A PET study. Aging Brain. 2022;2:100048. doi:10.1016/j.nbas.2022.100048

43. Frings L, Hellwig S, Spehl TS, et al. Asymmetries of amyloid-β burden and neuronal dysfunction are positively correlated in Alzheimer’s disease. Brain. 2015;138(10):3089–3099. doi:10.1093/brain/awv229

44. Martersteck A, Murphy C, Rademaker A, et al. Is in vivo amyloid distribution asymmetric in primary progressive aphasia? Ann Neurol. 2016;79(3):496–501. doi:10.1002/ana.24566

45. Lehmann M, Ghosh PM, Madison C, et al. Diverging patterns of amyloid deposition and hypometabolism in clinical variants of probable Alzheimer’s disease. Brain J Neurol. 2013;136(Pt 3):844–858. doi:10.1093/brain/aws327

46. Lam AD, Thibault EG, Mayblyum DV, et al. Association of Seizure Foci and Location of Tau and Amyloid Deposition and Brain Atrophy in Patients With Alzheimer Disease and Seizures. Neurology. 2024;103(9):e209920. doi:10.1212/WNL.0000000000209920

47. Charil A, Cao Y, Willis BA, Hersch S, Irizarry MC, Reyderman L. Lecanemab Slows Tau PET Accumulation. Alzheimers Dement. 2024;20(S6):e091956. doi:10.1002/alz.091956

48. Milloz A, Vogel J, Olsen A, et al. Multiscale Quantification of Hemispheric Asymmetry in Cortical Maps Using Geometric Eigenmodes. Published online October 31, 2024:2024.10.31.621232. doi:10.1101/2024.10.31.621232

49. Sims JR, Zimmer JA, Evans CD, et al. Donanemab in Early Symptomatic Alzheimer Disease: The TRAILBLAZER-ALZ 2 Randomized Clinical Trial. JAMA. 2023;330(6):512–527. doi:10.1001/jama.2023.13239

50. Budd Haeberlein S, Aisen PS, Barkhof F, et al. Two Randomized Phase 3 Studies of Aducanumab in Early Alzheimer’s Disease. J Prev Alzheimers Dis. 2022;9(2):197–210. doi:10.14283/jpad.2022.30

51. Dyck CH van, Swanson CJ, Aisen P, et al. Lecanemab in Early Alzheimer’s Disease. N Engl J Med. 2023;388(1):9–21. doi:10.1056/NEJMoa2212948

52. Brum WS, Cullen NC, Janelidze S, et al. A two-step workflow based on plasma p-tau217 to screen for amyloid β positivity with further confirmatory testing only in uncertain cases. Nat Aging. 2023;3(9):1079–1090. doi:10.1038/s43587-023-00471-5

53. Gobom J, Parnetti L, Rosa-Neto P, et al. Validation of the LUMIPULSE automated immunoassay for the measurement of core AD biomarkers in cerebrospinal fluid. Clin Chem Lab Med CCLM. 2022;60(2):207–219. doi:10.1515/cclm-2021-0651

54. Coomans EM, Tomassen J, Ossenkoppele R, et al. Genetically identical twins show comparable tau PET load and spatial distribution. Brain. 2022;145(10):3571–3581. doi:10.1093/brain/awac004

55. Pollard TJ, Johnson AEW, Raffa JD, Mark RG. tableone: An open source Python package for producing summary statistics for research papers. JAMIA Open. 2018;1(1):26–31. doi:10.1093/jamiaopen/ooy012

56. Seabold S, Perktold J. Statsmodels: Econometric and Statistical Modeling with Python. In: ; 2010:92–96. doi:10.25080/Majora-92bf1922-011

57. Cho SH, Choe YS, Park S, et al. Appropriate reference region selection of 18F-florbetaben and 18F-flutemetamol beta-amyloid PET expressed in Centiloid. Sci Rep. 2020;10(1):14950. doi:10.1038/s41598-020-70978-z

58. Baker SL, Maass A, Jagust WJ. Considerations and code for partial volume correcting [18F]-AV-1451 tau PET data. Data Brief. 2017;15:648–657. doi:10.1016/j.dib.2017.10.024

59. Thurfjell L, Lilja J, Lundqvist R, et al. Automated quantification of 18F-flutemetamol PET activity for categorizing scans as negative or positive for brain amyloid: concordance with visual image reads. J Nucl Med Off Publ Soc Nucl Med. 2014;55(10):1623–1628. doi:10.2967/jnumed.114.142109

60. Mattsson N, Palmqvist S, Stomrud E, Vogel J, Hansson O. Staging β-Amyloid Pathology With Amyloid Positron Emission Tomography. JAMA Neurol. 2019;76(11):1319–1329. doi:10.1001/jamaneurol.2019.2214

61. Ahmadi K, Pereira JB, Berron D, et al. Gray matter hypoperfusion is a late pathological event in the course of Alzheimer’s disease. J Cereb Blood Flow Metab Off J Int Soc Cereb Blood Flow Metab. 2023;43(4):565–580. doi:10.1177/0271678X221141139

62. Fischl B. FreeSurfer. NeuroImage. 2012;62(2):774-781. doi:10.1016/j.neuroimage.2012.01.021

63. Jenkinson M, Beckmann CF, Behrens TEJ, Woolrich MW, Smith SM. FSL. NeuroImage. 2012;62(2):782-790. doi:10.1016/j.neuroimage.2011.09.015

64. Tournier JD, Smith R, Raffelt D, et al. MRtrix3: A fast, flexible and open software framework for medical image processing and visualisation. NeuroImage. 2019;202:116137. doi:10.1016/j.neuroimage.2019.116137

65. Jon Clucas, Steve Giavasis, Amy Gutierrez, et al. FCP-INDI/C-PAC: C-PAC Version 1.3.0.post2 Beta. Published online May 14, 2024. doi:10.5281/ZENODO.593145

66. Abraham A, Pedregosa F, Eickenberg M, et al. Machine learning for neuroimaging with scikit-learn. Front Neuroinformatics. 2014;8. doi:10.3389/fninf.2014.00014

67. Esteban O, Markiewicz CJ, Burns C, et al. nipy/nipype: 1.8.3. Published online July 14, 2022. doi:10.5281/zenodo.6834519

68. Friston KJ. Functional and effective connectivity: a review. Brain Connect. 2011;1(1):13–36. doi:10.1089/brain.2011.0008

69. Desikan RS, Ségonne F, Fischl B, et al. An automated labeling system for subdividing the human cerebral cortex on MRI scans into gyral based regions of interest. NeuroImage. 2006;31(3):968–980. doi:10.1016/j.neuroimage.2006.01.021

70. Fischl B, Salat DH, Busa E, et al. Whole Brain Segmentation: Automated Labeling of Neuroanatomical Structures in the Human Brain. Neuron. 2002;33(3):341–355. doi:10.1016/S0896-6273(02)00569-X

71. Dhollander T, Mito R, Raffelt D, Connelly A. Improved white matter response function estimation for 3-tissue constrained spherical deconvolution. In: Proc Intl Soc Mag Reson Med, 2019, 555. ; 2019.

72. Jeurissen B, Tournier JD, Dhollander T, Connelly A, Sijbers J. Multi-tissue constrained spherical deconvolution for improved analysis of multi-shell diffusion MRI data. NeuroImage. 2014;103:411–426. doi:10.1016/j.neuroimage.2014.07.061

73. Smith R, Skoch A, Bajada C, Caspers S, Connelly A. Hybrid Surface-Volume Segmentation for improved Anatomically-Constrained Tractography. In: Proceedings of the Oganisation for Human Brain Mapping. ; 2020.

74. Tournier JD, Calamante F, Connelly A. Improved probabilistic streamlines tractography by 2nd order integration over fibre orientation distributions. In: Proceedings of the International Society for Magnetic Resonance in Medicine, 2010, 1670. ; 2010.

75. Smith RE, Tournier JD, Calamante F, Connelly A. Anatomically-constrained tractography: Improved diffusion MRI streamlines tractography through effective use of anatomical information. NeuroImage. 2012;62(3):1924–1938. doi:10.1016/j.neuroimage.2012.06.005

76. Smith RE, Tournier JD, Calamante F, Connelly A. SIFT2: Enabling dense quantitative assessment of brain white matter connectivity using streamlines tractography. NeuroImage. 2015;119:338–351. doi:10.1016/j.neuroimage.2015.06.092

77. Basser PJ, Mattiello J, Lebihan D. Estimation of the Effective Self-Diffusion *Tensor* from the NMR Spin Echo. J Magn Reson B. 1994;103(3):247–254. doi:10.1006/jmrb.1994.1037

78. Basser PJ, Mattiello J, LeBihan D. MR diffusion tensor spectroscopy and imaging. Biophys J. 1994;66(1):259–267. doi:10.1016/S0006-3495(94)80775-1

79. Wasserthal J, Neher P, Maier-Hein KH. TractSeg - Fast and accurate white matter tract segmentation. NeuroImage. 2018;183:239–253. doi:10.1016/j.neuroimage.2018.07.070

80. Zalesky A, Fornito A, Bullmore ET. Network-based statistic: Identifying differences in brain networks. NeuroImage. 2010;53(4):1197–1207. doi:10.1016/j.neuroimage.2010.06.041

81. Schreiber S, Landau SM, Fero A, Schreiber F, Jagust WJ, for the Alzheimer’s Disease Neuroimaging Initiative. Comparison of Visual and Quantitative Florbetapir F 18 Positron Emission Tomography Analysis in Predicting Mild Cognitive Impairment Outcomes. JAMA Neurol. 2015;72(10):1183-1190. doi:10.1001/jamaneurol.2015.1633

82. Ossenkoppele R, Leuzy A, Cho H, et al. The impact of demographic, clinical, genetic, and imaging variables on tau PET status. Eur J Nucl Med Mol Imaging. 2021;48(7):2245–2258. doi:10.1007/s00259-020-05099-w

