## Supplementary Info for "Hemispheric Asymmetry of Tau Pathology is Related to Asymmetric Amyloid Deposition in Alzheimer’s Disease"

### Supplementary S1 – Extended methods

#### Inclusion and exclusion criteria for the participants

The Swedish BioFINDER-2 study enrolls participants in five sub-cohorts. Cohort A and B includes neurologically and cognitively healthy controls. The inclusion criteria are: i) ages 40-65 years (cohort A) and ages 66-100 years (cohort B); ii) absence of cognitive symptoms as assessed by a physician with special interest in cognitive disorders; iii) Mini Mental State Examination (MMSE) score 27-30 (cohort A) or 26-30 (cohort B) at screening visit; iv) do not fulfill the criteria for MCI or any dementia according to DSM-5[1]; v) fluent in Swedish.

Cohort C comprises participants with subjective cognitive deficits (SCD), or mild cognitive impairment (MCI; defined as a performance of <-1.5 SD below reference mean in at least one cognitive domain, see [2] for further details). Inclusion criteria are: i) Age 40-100 years; ii) referred to the memory clinics due to cognitive symptoms; iii) MMSE score of 24 – 30 points; iv) does not fulfill the criteria for any dementia (major neurocognitive disorder) according to DSM-5, v) fluent in Swedish.

Cohort D consists of participants with dementia due to AD. Inclusion criteria are: i) Age 40-100 years; ii) referred to the memory clinics due to cognitive symptoms; iii) MMSE score of > 12 points; iv) fulfill the DSM-5 criteria for dementia (major neurocognitive disorder) due to Alzheimer's disease[1]; v) fluent in Swedish.

Cohort E covers other non-AD dementias and neurodegenerative disorders. Inclusion criteria are: i) Age 40-100 years; ii) fulfillment of criteria for dementia (major neurocognitive disorder) due to Frototemporal dementia (FTD), Parkinson's disease with dementia (PDD), dementia with Lewy bodies (DLB) or subcortical VaD accordingly to the DSM-5 alternatively the criteria for Parkinson's disease (PD),[3] progressive supranuclear palsy (PSP),[4] multiple system atrophy (MSA),[5] corticobasal syndrome (CBS)[6] or semantic variant primary progressive aphasia (svPPA)[7]; iii) fluent in Swedish. Exclusion criteria for all sub-cohorts are: i) significant unstable systemic illness that makes it difficult to participate in the study; ii) current significant alcohol or substance misuse; iii) refusing lumbar puncture, MRI or PET.

The participants in the present study had been enrolled in either cohort A, B, or C (only people with SCD) of the BioFINDER-2 study for the cognitively unimpaired (CU) group, in cohort C for the MCI group, or D for the AD group. All participants were assessed by physicians with expertise in dementia disorders.

### Regions of interest

The meta regions of interest (meta-ROIs) included known regions relevant to neuropathological progression in neurodegenerative diseases, such as Braak staging for tau pathology and Aβ staging based on its progression.[8–10] The meta-ROIs also included commonly used temporal meta-ROI which is based on Braak I-IV and global (i.e., whole-brain) meta-ROI. See Table S1.1 for detailed overview of the meta-ROIs and the regions involved.

***Table S1.1***. *Meta regions of interest.*

| Meta-ROI | Regions involved |
|---|---|
| **Global** | *whole brain* (i.e., all Desikan-Killiany regions) |
| **Temporal** | entorhinal cortex, parahippocampal cortex, fusiform cortex, amygdala, inferior temporal cortex, middle temporal cortex |
| **Braak I-II** | entorhinal cortex |
| **Braak III-IV** | parahippocampal cortex, fusiform cortex, amygdala, inferior temporal cortex, middle temporal cortex |
| **Braak V-VI** | caudal anterior cingulate cortex, caudal middle frontal cortex, cuneus, inferior parietal cortex, isthmus cingulate cortex, lateral occipital cortex, lateral orbitofrontal cortex, lingual cortex, medial orbitofrontal cortex, paracentral cortex, pars opercularis, pars triangularis, pars orbitalis, pericalcarine cortex, postcentral cortex, posterior cingulate cortex, precentral cortex, precuneus, rostral anterior cingulate cortex, rostral middle frontal cortex, superior frontal cortex, superior parietal cortex, superior temporal cortex, supramarginal cortex, frontal pole, temporal pole, transverse temporal cortex, insula |

| | |
|---|---|
| **Early-Aβ** | precuneus, posterior cingulate cortex, isthmus cingulate cortex, insula, medial orbitofrontal cortex, lateral orbitofrontal cortex |
| **Intermediate-Aβ** | banks of superior temporal sulcus, caudal middle frontal cortex, cuneus, frontal pole, fusiform cortex, inferior parietal cortex, inferior temporal cortex, lateral occipital cortex, middle temporal cortex, parahippocampal cortex, pars opercularis, pars orbitalis, pars triangularis, putamen, rostral anterior cingulate cortex, rostral middle frontal cortex, superior frontal cortex, superior parietal cortex, superior temporal cortex, supramarginal cortex |
| **Late-Aβ** | lingual cortex, pericalcarine cortex, paracentral cortex, precentral cortex, postcentral cortex |

#### Structural connectivity, microstructural integrity, and functional connectivity

The estimation of structural connectivity (SC) was performed using NiPype,[11] Mrtrix3,[12] FSL[13] and FreeSurfer[14] software packages. Firstly, a single set of response functions for white matter (WM), gray matter (GM) and cerebrospinal fluid (CSF) were estimated by "dhollander" algorithm[15,16] using pre-processed dMRI data from 60 CU A-T- and 40 CU A+T- participants of the BioFinder2 (BF2) cohort. These were then utilised to estimate fiber orientation distributions (FOD) based on multi-shell multi-tissue Constrained Spherical Deconvolution (CSD)[17] using "msmt_csd" algorithm,[18] which enables to calculate three separate FODs for three tissue types (i.e., WM, GM, CSF) based on multi-shell dMRI data. Second, a five-tissue-type (5TT) segmented tissue image was generated based on Hybrid Surface and Volume Segmentation (HSVS) using "hsvs" algorithm[19] which uses FreeSurfer and FSL to create segmentations of different tissue types (i.e, cortical GM, sub-cortical GM, WM, CSF, and optionally pathological tissue). 5TT image is important for employing anatomical constraints for later fiber tracking i.e. it increases biological plausibility of the tractogram. After that, the 5TT and T1-weighted images are co-registered to the dMRI data using FSL's "flirt"[20,21] with the average of the not gradient weighted b0 dMRI data. Third, Anatomically-Constrained Tractography (ACT) was performed using a probabilistic "iFOD2" tracking algorithm[22,23] by estimating 10 million streamlines with applying the 5TT image and dynamic determination of seed points.[24] Following that, to reduce the bias in overestimation of streamlines compared to biological WM fibres, Spherical-deconvolution Informed Filtering of Tractograms 2 (SIFT2) method[24] was utilised to calculate weights for all streamlines. Structural connectivity (SC) matrix was then generated based on Desikan-Killiany atlas[25] and ASEG protocol[26] (i.e., 84 regions from FreeSurfer's "aparcaseg") using the sum of SIFT2-weighted streamlines as weights of the edges.[27]

For estimating microstructural integrity, diffusion tensor imaging (DTI) was applied to the dMRI data using weighted least-squares method[28] and removing the b2500 volumes from dMRI data beforehand (i.e., using only b0, b100, b1000 shells). Then, maps of fractional anisotropy (FA) and mean diffusivity (MD) were calculated from the tensor model.[29] White matter tract segmentation was performed using TractSeg, a convolutional neural network-based approach that directly segments tracts from fiber orientation distribution function peaks.[30] The algorithm generated bundle segmentations of the main inter-hemispheric white matter tracts (corpus callosum, forceps major, forceps minor), from which mean FA and MD values were extracted for each tract.

Functional connectivity (FC) was estimated using Nilearn software.[31] FC were constructed from the subject-space pre-processed resting state fMRI data by extracting time series data[32] from the same Desikan-Killiany regions as previously done for SC. After that, Pearson correlation with Fisher's z-transformation[33] was applied between all brain regions for calculating FC.

### Supplementary S2 – Extended results

#### Association between tau laterality index and continuous measure of inter-hemispheric connectivity

To ensure that our null finding regarding the association between inter-hemispheric connectivity and tau asymmetry was not due to the grouping definitions, we performed an additional analysis using a linear regression with continuous tau laterality index instead of the group comparison. Similarly, this analysis resulted in no association between tau laterality and inter-hemispheric functional connectivity (β=0.090, p=0.130) or structural connectivity (β=-0.037, p=0.502) at Global meta-ROI (Fig. S2.1).

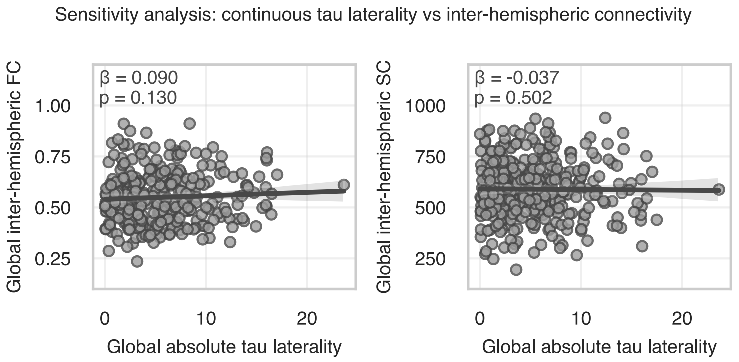


***Figure S2.1****. Association between absolute tau laterality index and inter-hemispheric FC (left) and SC (right). FC – functional connectivity; SC – structural connectivity*

#### Mean diffusivity across the main tracts connecting the two hemispheres

Similarly to the analysis of comparing fractional anisotropy of the three white matter tracts between the tau asymmetry groups, mean diffusivity revealed no significant differences between the subjects displaying asymmetrical tau distribution compared to individuals with symmetric tau (Fig. S2.2) – in the corpus callosum (S-LA: β=-0.065, 95%CI=[-0.311; 0.181], p>0.9; S-RA: β=0.040, 95%CI=[-0.329; 0.408], p>0.9), forceps major (S-LA: β=-0.013, 95%CI=[-0.264; 0.238], p>0.9; S-RA: β=0.210, 95%CI=[-0.159; 0.578], p=0.790), or forceps minor (S-LA: β=0.112, 95%CI=[-0.137; 0.361], p>0.9; S-RA: β=0.148, 95%CI=[-0.223; 0.520], p>0.9).

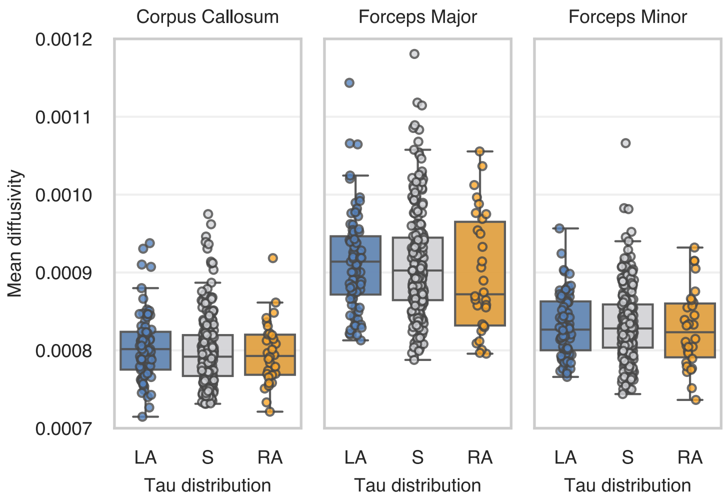


***Figure S2.2***. *Average mean diffusivity across three main inter-hemispheric white matter tracts. LA – left tau asymmetric; S – tau symmetric – RA – right tau asymmetric.*

#### Network Based Statistics

Whole-brain connectome analysis using Network Based Statistics (NBS) showed some minor differences between the tau asymmetry groups (Table S2.1; Fig. S2.2). Higher functional connectivity in the left asymmetric group compared to the symmetric group was found in a cluster of nodes primarily located in the right hemisphere which also included a few inter-hemispheric connections (threshold=3.0; C=9, p=0.018). The right asymmetric group showed reduced structural connectivity compared to the symmetric group in a cluster of nodes within the right hemisphere (threshold=3.0; C=4, p=0.016). However, the results were not consistent across statistical threshold. Therefore, we considered these results not entirely reliable. Moreover, these differences are likely due to the unbalanced tau burden in the affected hemisphere for subjects with asymmetric tau distribution which we were not able properly adjust for at the single node level in NBS (i.e., we adjusted for global average tau uptake).

***Table S2.1***. *NBS analysis comparisons of functional and structural connectivity between tau asymmetry groups. Each column represents a contrast between groups and each row shows the t-statistic threshold (t) used. Each cell represents the maximum detected component size C and the significance level i.e., C (p-value). A – tau* *asymmetric; LA – left tau asymmetric; S – tau symmetric – RA – right tau asymmetric; FC – functional connectivity; SC – structural connectivity.*

| **FC** | **A > S** | **A < S** | **LA > S** | **LA < S** | **RA > S** | **RA < S** |
|---|---|---|---|---|---|---|
| **t = 2.5** | 19 (p=0.045) | - | 23 (p=0.041) | - | 11 (p=0.072) | 2 (p=0.239) |
| **t = 3.0** | 2 (p=0.077) | - | 9 (p=0.018) | - | 3 (p=0.056) | 1 (p=0.161) |
| **t = 3.5** | - | - | 1 (p=0.053) | - | 1 (p=0.054) | - |
| **SC** | **A > S** | **A < S** | **LA > S** | **LA < S** | **RA > S** | **RA < S** |
| **t = 2.5** | 1 (p=0.684) | 5 (p=0.104) | 3 (p=0.231) | 6 (p=0.074) | 2 (p=0.423) | 13 (p=0.031) |
| **t = 3.0** | - | 2 (p=0.081) | 1 (p=0.384) | 1 (p=0.241) | 2 (p=0.156) | 4 (p=0.016) |
| **t = 3.5** | - | 1 (p=0.073) | 1 (p=0.127) | - | - | - |

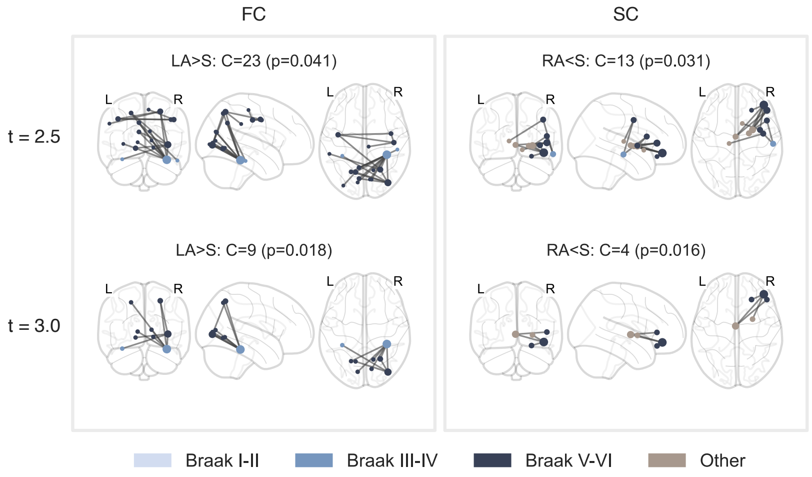


***Figure S2.3**. Components detected using NBS analysis between the tau asymmetry groups. FC – functional connectivity; SC – structural connectivity; LA – left tau asymmetric; S – tau symmetric – RA – right tau asymmetric; C – component size; NBS – Network Based Statistic; t – threshold used in NBS*

#### Associations between the laterality of Aβ and tau across different meta-ROIs

Between tau laterality and Aβ laterality, an additional set of analyses was performed using different meta-ROIs than used in the main analysis. First, the relationship between global tau laterality and Aβ laterality at Aβ staging meta-ROIs was assessed (Fig. S2.4a), which resulted the strongest effect size at Intermediate-Aβ meta-ROI (β=0.634, 95%CI=[0.533; 0.735], p<0.001) followed by Late-Aβ (β=0.609, 95%CI=[0.505; 0.713], p<0.001) and Early-Aβ (β=0.547, 95%CI=[0.438; 0.657], p<0.001). Second, the association between global Aβ laterality and tau laterality at different Braak stages was investigated (Fig. S2.4b), with the strongest effect found at Braak III-IV meta-ROI (β=0.654, 95%CI=[0.555; 0.753], p<0.001) followed by Braak V-VI (β=0.594, 95%CI=[0.488; 0.699], p<0.001) and Braak I-II (β=0.443, 95%CI=[0.327; 0.559], p<0.001).

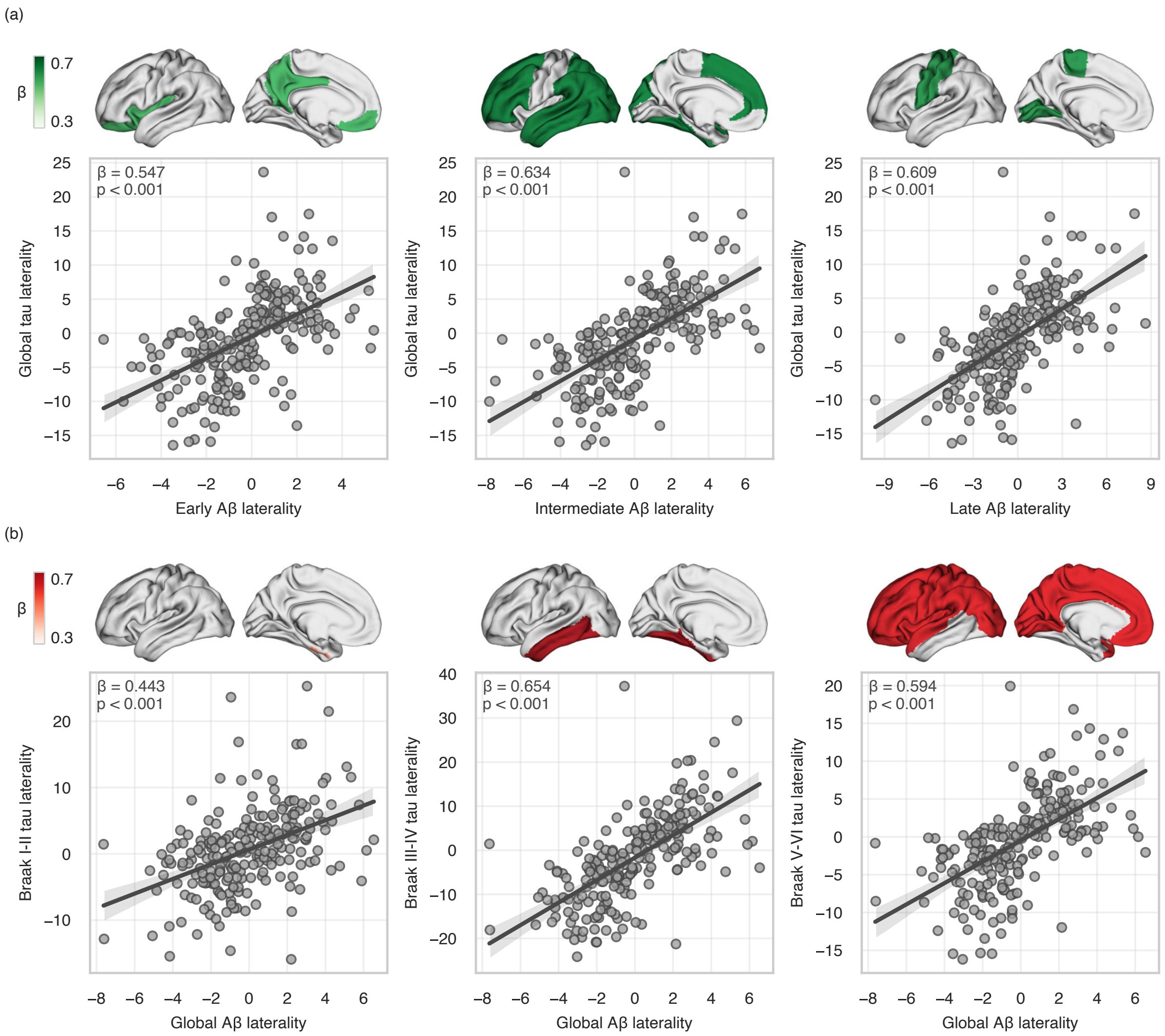


***Figure S2.4***. *Associations between Aβ laterality and tau laterality across meta-ROIs: (a) Global tau laterality vs Aβ laterality at Aβ stages; (b) Global Aβ laterality vs tau laterality at Braak stages.*
*Aβ – amyloid-beta*

#### Difference in estimated Aβ onset between hemispheres between the groups

Time difference in Aβ onset between hemispheres was computed using the sampled iterative local approximation (SILA) algorithm[34] to investigate inter-hemispheric difference in Aβ accumulation across the groups defined based on tau laterality index. The SILA algorithm was applied to the full longitudinal BioFINDER-2 cohort to model estimated years until/since Aβ onset for each hemisphere for each meta-ROI. These unilateral values were subtracted within each meta-ROI to obtain the estimated time difference of Aβ onset between hemispheres. For the comparison of the estimated hemispheric time difference in Aβ onset between the groups, OLS multiple linear regressions (OLS: estimated Aβ onset difference ~ age + sex + group) was used with the significance level set to $p < 0.05$ after Bonferroni correction accounting for the multiple comparisons between groups (i.e., $\text{p-val}_{\text{Bonf.}} = \text{p-val} * 3$).

We found that the difference in estimated onset of global Aβ pathology between hemispheres was significantly higher in the asymmetric groups compared to the symmetric group (Fig. 4c) with a median difference in the left asymmetric of 1.7 years (t=3.123, p=0.006) and of 2.5 years in the right asymmetric (t=4.137, p<0.001), compared to 1.3 years in the symmetric group, which was similar across other meta-ROIs (Fig. S2.5).

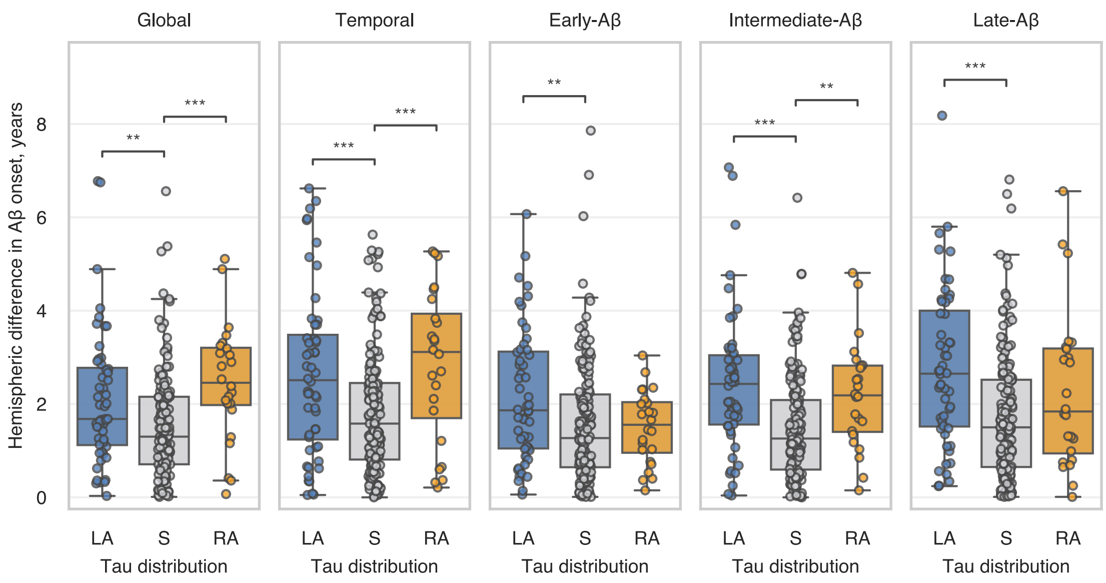


***Figure S2.5***. *Comparison of the SILA-based estimation of time difference in Aβ onset between hemispheres across meta-ROIs.*
*Aβ – amyloid-beta; SILA – sampled iterative local approximation algorithm; LA – left tau asymmetric; S – tau symmetric – RA – right tau asymmetric; ** – p<0.01; *** – p<0.001.*

### Summary of the longitudinal sample

***Table S2.2***. *Demographics of the longitudinal dataset.*
*Categorical variables have been presented as 'count (%)', normally distributed continuous variables as 'mean (SD)' and non-normally distributed variables as 'median [IQR]'. T- – tau negative; T+ – tau positive; M – male; F – female; CU – cognitively unimpaired; MCI – mild cognitive impairment; AD – Alzheimer's disease; SUVR – standardised uptake value ratio; LI – laterality index; Aβ – amyloid-beta; MMSE – Mini-Mental State Examination; mPACC – modified Preclinical Alzheimer Cognitive Composite.*

| | | **Longitudinal A+ (n=289)** | | |
|---|---|---|---|---|
| | | **T- (n=180)** | **T+ (n=109)** | **P-value** |
| **Age, years** | | 73.09 (7.89) | 72.08 (7.06) | 0.264 |
| **Sex** | **M** | 92 (51%) | 57 (52%) | 0.941 |
| | **F** | 88 (49%) | 52 (48%) | |
| **Education, years** | | 12.78 (4.08) | 12.98 (3.68) | 0.670 |
| **Diagnosis** | **CU** | 116 (64%) | 36 (33%) | <0.001 |
| | **MCI** | 64 (36%) | 69 (63%) | |
| | **AD** | 0 (0%) | 4 (4%) | |
| **Temporal tau, SUVR** | | 1.19 [1.14,1.24] | 1.61 [1.42,2.00] | <0.001 |
| **Absolute Temporal tau LI** | | 1.31 (1.17) | 8.32 (6.10) | <0.001 |
| **Global Aβ, SUVR** | | 1.27 (0.19) | 1.50 (0.20) | <0.001 |
| **ApoE4** | **0** | 62 (34%) | 17 (16%) | 0.001 |
| | **1** | 99 (55%) | 71 (65%) | |
| | **2** | 19 (11%) | 21 (19%) | |
| **MMSE** | | 29.00 [27.00,29.25] | 27.00 [26.00,29.00] | <0.001 |

| mPACC | -0.56 [-1.53,-0.00] | -1.50 [-2.40,-0.63] | <0.001 |
|---|---|---|---|

#### Longitudinal association between baseline Aβ laterality and changes over time in tau laterality

***Table S2.3***. *Summary of the linear mixed effects model within A+ subsample predicting changes over time in tau laterality with baseline Aβ laterality using the Global meta-ROI.*
*The statistical significance of the baseline Aβ laterality and its interaction with time are annotated with* ***green*** *if it reached p<0.05 and* ***red*** *if it did not.*

| A+ (n=289 with a total of 707 datapoints)<br>LME: Tau LI ~ time * ($Age_{BAS}$ + Sex + Aβ $LI_{BAS}$) + [time \| participant] | | | | |
|---|---|---|---|---|
| **ROI** | **Global** | | | |
| | **β** | **SE** | **95% CI** | **p** |
| **Intercept** | -0.052 | 0.056 | [-0.162, 0.058] | 0.350 |
| **Time** | -0.001 | 0.016 | [-0.032, 0.031] | 0.964 |
| **$Age_{BAS}$** | -0.007 | 0.040 | [-0.086, 0.072] | 0.864 |
| **Sex** | 0.023 | 0.081 | [-0.135, 0.181] | 0.776 |
| **Aβ $LI_{BAS}$** | 0.379 | 0.040 | [0.301, 0.458] | **<0.001** |
| **Time × $Age_{BAS}$** | -0.000 | 0.012 | [-0.023, 0.022] | 0.994 |
| **Time × Sex** | -0.000 | 0.023 | [-0.046, 0.045] | 0.995 |
| **Time × Aβ $LI_{BAS}$** | 0.025 | 0.012 | [0.003, 0.048] | **0.028** |

***Table S2.4***. *Summary of the linear mixed effects model within A+T- subsample predicting tau laterality over time with baseline Aβ laterality at Braak meta-ROIs.*
*The statistical significance of the baseline Aβ laterality and its interaction with time are annotated with* ***green*** *if it reached p<0.05 and* ***red*** *if it did not. Furthermore,* * *indicates whether the p-value survived the statistical threshold after Bonferroni correction.*

| A+T- (n=180 with a total of 452 datapoints)<br>LME: **Tau LI** ~ **time** * ($Age_{BAS}$ + Sex + **Aβ $LI_{BAS}$**) + [time \| participant] | | | | | | | | | | | | |
|---|---|---|---|---|---|---|---|---|---|---|---|---|
| **ROI** | **Braak I-II** | | | | **Braak III-IV** | | | | **Braak V-VI** | | | |
| | **β** | **SE** | **95% CI** | **p** | **β** | **SE** | **95% CI** | **p** | **β** | **SE** | **95% CI** | **p** |
| **Intercept** | -0.014 | 0.106 | [-0.221, 0.193] | 0.894 | -0.041 | 0.067 | [-0.173, 0.092] | 0.546 | -0.046 | 0.076 | [-0.195, 0.102] | 0.539 |
| **Time** | 0.030 | 0.026 | [-0.021, 0.082] | 0.250 | 0.021 | 0.029 | [-0.036, 0.078] | 0.479 | 0.013 | 0.035 | [-0.056, 0.081] | 0.713 |
| **$Age_{BAS}$** | -0.013 | 0.075 | [-0.161, 0.134] | 0.861 | -0.044 | 0.048 | [-0.138, 0.050] | 0.360 | -0.096 | 0.054 | [-0.201, 0.010] | 0.075 |
| **Sex** | -0.023 | 0.151 | [-0.318, 0.273] | 0.881 | -0.086 | 0.096 | [-0.274, 0.103] | 0.372 | 0.017 | 0.107 | [-0.193, 0.228] | 0.871 |

| | | | | | | | | | | | | |
|---|---|---|---|---|---|---|---|---|---|---|---|---|
| **Aβ $LI_{BAS}$** | 0.067 | 0.074 | [-0.077, 0.212] | **0.363** | 0.246 | 0.048 | [0.152, 0.339] | **<0.001*** | 0.202 | 0.053 | [0.098, 0.306] | **<0.001*** |
| **Time × $Age_{BAS}$** | -0.026 | 0.019 | [-0.063, 0.010] | 0.158 | -0.019 | 0.021 | [-0.059, 0.022] | 0.366 | -0.018 | 0.025 | [-0.066, 0.030] | 0.464 |
| **Time × Sex** | -0.041 | 0.037 | [-0.113, 0.032] | 0.269 | 0.059 | 0.041 | [-0.022, 0.139] | 0.153 | 0.036 | 0.049 | [-0.060, 0.131] | 0.464 |
| **Time × Aβ $LI_{BAS}$** | 0.054 | 0.019 | [0.017, 0.091] | **0.004*** | 0.080 | 0.020 | [0.039, 0.120] | **<0.001*** | 0.048 | 0.025 | [-0.000, 0.096] | **0.050** |

***Table S2.5***. *Summary of the linear mixed effects model within A+T+ subsample predicting tau laterality over time with baseline Aβ laterality at Braak meta-ROIs.*
*The statistical significance of the baseline Aβ laterality and its interaction with time are annotated with **green** if it reached p<0.05 and **red** if it did not. Furthermore, * indicates whether the p-value survived the statistical threshold after Bonferroni correction.*

A+T+ (n=109 with a total of 255 datapoints)

LME: **Tau LI** ~ **time** * ($Age_{BAS}$ + Sex + **Aβ $LI_{BAS}$**) + [time | participant]

| **ROI** | **Braak I-II** | | | | **Braak III-IV** | | | | **Braak V-VI** | | | |
|---|---|---|---|---|---|---|---|---|---|---|---|---|
| | **β** | **SE** | **95% CI** | **p** | **β** | **SE** | **95% CI** | **p** | **β** | **SE** | **95% CI** | **p** |
| **Intercept** | -0.047 | 0.121 | [-0.284, 0.190] | 0.700 | -0.088 | 0.075 | [-0.235, 0.059] | 0.242 | -0.054 | 0.074 | [-0.199, 0.091] | 0.466 |
| **Time** | -0.010 | 0.036 | [-0.080, 0.060] | 0.778 | 0.002 | 0.020 | [-0.037, 0.041] | 0.917 | -0.017 | 0.027 | [-0.069, 0.035] | 0.526 |
| **$Age_{BAS}$** | -0.058 | 0.089 | [-0.232, 0.116] | 0.511 | -0.032 | 0.055 | [-0.140, 0.075] | 0.555 | 0.015 | 0.054 | [-0.091, 0.122] | 0.781 |
| **Sex** | 0.041 | 0.175 | [-0.303, 0.384] | 0.817 | 0.108 | 0.109 | [-0.106, 0.323] | 0.321 | 0.068 | 0.107 | [-0.143, 0.278] | 0.528 |
| **Aβ $LI_{BAS}$** | 0.280 | 0.088 | [0.107, 0.453] | **0.002*** | 0.679 | 0.054 | [0.573, 0.784] | **<0.001*** | 0.584 | 0.053 | [0.481, 0.688] | **<0.001*** |
| **Time × $Age_{BAS}$** | 0.009 | 0.026 | [-0.042, 0.061] | 0.719 | 0.002 | 0.015 | [-0.027, 0.030] | 0.898 | 0.003 | 0.020 | [-0.035, 0.042] | 0.871 |
| **Time × Sex** | 0.063 | 0.054 | [-0.043, 0.169] | 0.243 | -0.008 | 0.030 | [-0.068, 0.051] | 0.781 | -0.009 | 0.039 | [-0.087, 0.068] | 0.810 |
| **Time × Aβ $LI_{BAS}$** | 0.024 | 0.026 | [-0.028, 0.076] | **0.369** | 0.004 | 0.015 | [-0.026, 0.034] | **0.783** | 0.033 | 0.019 | [-0.005, 0.071] | **0.090** |

***Table S2.6***. *Summary of the linear mixed effects model within A+T- subsample who stay A+T- throughout follow-up predicting tau laterality over time with baseline Aβ laterality at Braak meta-ROIs.*
*The statistical significance of the baseline Aβ laterality and its interaction with time are annotated with **green** if it reached p<0.05 and **red** if it did not. Furthermore, * indicates whether the p-value survived the statistical threshold after Bonferroni correction.*

A+T- to A+T- (n=142 with a total of 347 datapoints)

LME: **Tau LI** ~ **time** * ($Age_{BAS}$ + Sex + **Aβ $LI_{BAS}$**) + [time | participant]

| **ROI** | **Braak I-II** | | | | **Braak III-IV** | | | | **Braak V-VI** | | | |
|---|---|---|---|---|---|---|---|---|---|---|---|---|
| | **β** | **SE** | **95% CI** | **p** | **β** | **SE** | **95% CI** | **p** | **β** | **SE** | **95% CI** | **p** |

| | | | | | | | | | | | | |
|---|---|---|---|---|---|---|---|---|---|---|---|---|
| **Intercept** | 0.010 | 0.121 | [-0.227, 0.246] | 0.936 | 0.009 | 0.094 | [-0.176, 0.193] | 0.924 | -0.048 | 0.097 | [-0.238, 0.141] | 0.616 |
| **Time** | 0.036 | 0.030 | [-0.023, 0.096] | 0.230 | 0.039 | 0.039 | [-0.038, 0.116] | 0.320 | 0.034 | 0.038 | [-0.039, 0.108] | 0.363 |
| **Age$_{BAS}$** | 0.014 | 0.087 | [-0.157, 0.185] | 0.872 | -0.066 | 0.068 | [-0.199, 0.067] | 0.332 | -0.117 | 0.070 | [-0.254, 0.020] | 0.093 |
| **Sex** | -0.055 | 0.174 | [-0.396, 0.287] | 0.753 | -0.107 | 0.136 | [-0.372, 0.159] | 0.431 | 0.001 | 0.139 | [-0.272, 0.274] | 0.995 |
| **Aβ LI$_{BAS}$** | 0.039 | 0.086 | [-0.129, 0.206] | **0.651** | 0.268 | 0.067 | [0.137, 0.400] | **<0.001*** | 0.171 | 0.069 | [0.035, 0.307] | **0.013*** |
| **Time × Age$_{BAS}$** | -0.014 | 0.022 | [-0.056, 0.029] | 0.529 | 0.024 | 0.028 | [-0.031, 0.079] | 0.386 | 0.015 | 0.026 | [-0.037, 0.067] | 0.568 |
| **Time × Sex** | -0.045 | 0.043 | [-0.129, 0.039] | 0.298 | 0.006 | 0.056 | [-0.103, 0.116] | 0.911 | 0.007 | 0.053 | [-0.096, 0.109] | 0.902 |
| **Time × Aβ LI$_{BAS}$** | 0.076 | 0.022 | [0.033, 0.119] | **0.001*** | 0.035 | 0.028 | [-0.019, 0.089] | **0.202** | -0.001 | 0.026 | [-0.051, 0.050] | **0.979** |

***Table S2.7**. Summary of the linear mixed effects model within A+T- subsample who progress to A+T+ during follow-up predicting tau laterality over time with baseline Aβ laterality at Braak meta-ROIs. The statistical significance of the baseline Aβ laterality and its interaction with time are annotated with **green** if it reached p<0.05 and **red** if it did not. Furthermore, * indicates whether the p-value survived the statistical threshold after Bonferroni correction.*

A+T- to A+T+ (n=38 with a total of 105 datapoints)

LME: **Tau LI** ~ **time** * (Age$_{BAS}$ + Sex + **Aβ LI$_{BAS}$**) + [time | participant]

| **ROI** | **Braak I-II** | | | | **Braak III-IV** | | | | **Braak V-VI** | | | |
|---|---|---|---|---|---|---|---|---|---|---|---|---|
| | **β** | **SE** | **95% CI** | **p** | **β** | **SE** | **95% CI** | **p** | **β** | **SE** | **95% CI** | **p** |
| **Intercept** | -0.073 | 0.243 | [-0.550, 0.405] | 0.766 | -0.105 | 0.141 | [-0.381, 0.170] | 0.453 | -0.058 | 0.131 | [-0.315, 0.199] | 0.658 |
| **Time** | -0.021 | 0.057 | [-0.133, 0.092] | 0.721 | -0.013 | 0.040 | [-0.092, 0.066] | 0.744 | -0.048 | 0.061 | [-0.167, 0.070] | 0.424 |
| **Age$_{BAS}$** | -0.136 | 0.166 | [-0.461, 0.189] | 0.411 | 0.017 | 0.098 | [-0.176, 0.209] | 0.866 | -0.050 | 0.090 | [-0.226, 0.126] | 0.579 |
| **Sex** | 0.053 | 0.335 | [-0.604, 0.710] | 0.875 | -0.059 | 0.193 | [-0.438, 0.320] | 0.761 | 0.078 | 0.179 | [-0.273, 0.428] | 0.664 |
| **Aβ LI$_{BAS}$** | 0.184 | 0.166 | [-0.141, 0.509] | **0.268** | 0.326 | 0.099 | [0.132, 0.519] | **0.001*** | 0.303 | 0.087 | [0.132, 0.475] | **0.001*** |
| **Time × Age$_{BAS}$** | -0.077 | 0.038 | [-0.151, -0.003] | 0.041 | -0.092 | 0.028 | [-0.147, -0.037] | 0.001 | -0.082 | 0.042 | [-0.165, -0.000] | 0.050 |
| **Time × Sex** | 0.000 | 0.075 | [-0.147, 0.148] | 0.998 | 0.142 | 0.054 | [0.037, 0.247] | 0.008 | 0.099 | 0.082 | [-0.061, 0.260] | 0.224 |
| **Time × Aβ LI$_{BAS}$** | -0.003 | 0.037 | [-0.075, 0.070] | **0.942** | 0.132 | 0.028 | [0.077, 0.187] | **<0.001*** | 0.144 | 0.043 | [0.060, 0.227] | **0.001*** |

### Longitudinal association between baseline Aβ/tau laterality and changes over time in mPACC scores

***Table S2.8***. *Summary of the linear mixed effects models predicting mPACC scores over time with pathological asymmetry across the Braak meta-ROIs: (1) baseline tau laterality as predictor; (2) baseline tau laterality as predictor after adjusting for tau load; (3) baseline Aβ laterality as predictor.*
*The statistical significance of the interaction between time and baseline Aβ/tau laterality is annotated with* ***green*** *if it reached $p<0.05$ and* ***red*** *if it did not. Furthermore,* *** *indicates whether the p-value survived the statistical threshold after Bonferroni correction.*

| A+ (n=259 with a total of 606 datapoints) | | | | | | | | | | | | |
|---|---|---|---|---|---|---|---|---|---|---|---|---|
| LME #1: **mPACC** ~ **time** * ($Age_{BAS}$ + Sex + **Tau $LI_{BAS}$**) + [time \| participant] | | | | | | | | | | | | |
| **ROI** | **Braak I-II** | | | | **Braak III-IV** | | | | **Braak V-VI** | | | |
| | **β** | **SE** | **95% CI** | **p** | **β** | **SE** | **95% CI** | **p** | **β** | **SE** | **95% CI** | **p** |
| **Intercept** | 0.139 | 0.060 | [0.020, 0.257] | 0.022 | 0.140 | 0.057 | [0.028, 0.253] | 0.015 | 0.157 | 0.057 | [0.045, 0.270] | 0.006 |
| **Time** | -0.160 | 0.029 | [-0.216, -0.104] | <0.001 | -0.163 | 0.026 | [-0.214, -0.112] | <0.001 | -0.157 | 0.025 | [-0.205, -0.108] | <0.001 |
| **$Age_{BAS}$** | -0.122 | 0.043 | [-0.206, -0.038] | 0.004 | -0.129 | 0.041 | [-0.209, -0.050] | 0.001 | -0.147 | 0.041 | [-0.227, -0.067] | <0.001 |
| **Sex** | 0.170 | 0.086 | [0.002, 0.338] | 0.048 | 0.189 | 0.082 | [0.028, 0.349] | 0.021 | 0.153 | 0.082 | [-0.007, 0.313] | 0.060 |
| **Tau $LI_{BAS}$** | 0.034 | 0.043 | [-0.051, 0.119] | 0.432 | -0.203 | 0.040 | [-0.281, -0.126] | <0.001 | -0.206 | 0.040 | [-0.284, -0.128] | <0.001 |
| **Time × $Age_{BAS}$** | -0.015 | 0.021 | [-0.056, 0.025] | 0.462 | -0.021 | 0.019 | [-0.058, 0.016] | 0.256 | -0.035 | 0.018 | [-0.070, 0.000] | 0.053 |
| **Time × Sex** | -0.021 | 0.041 | [-0.101, 0.060] | 0.616 | -0.017 | 0.037 | [-0.089, 0.056] | 0.655 | -0.031 | 0.035 | [-0.100, 0.038] | 0.372 |
| **Time × Tau $LI_{BAS}$** | -0.036 | 0.020 | [-0.076, 0.004] | **0.076** | -0.134 | 0.019 | [-0.172, -0.095] | **<0.001 *** | -0.157 | 0.019 | [-0.194, -0.121] | **<0.001*** |
| LME #2: **mPACC** ~ **time** * ($Age_{BAS}$ + Sex + Tau $load_{BAS}$ + **Tau $LI_{BAS}$**) + [time \| participant] | | | | | | | | | | | | |
| **ROI** | **Braak I-II** | | | | **Braak III-IV** | | | | **Braak V-VI** | | | |
| | **β** | **SE** | **95% CI** | **p** | **β** | **SE** | **95% CI** | **p** | **β** | **SE** | **95% CI** | **p** |
| **Intercept** | 0.154 | 0.057 | [0.043, 0.266] | 0.007 | 0.157 | 0.056 | [0.047, 0.267] | 0.005 | 0.163 | 0.057 | [0.051, 0.275] | 0.004 |
| **Time** | -0.153 | 0.026 | [-0.204, -0.101] | <0.001 | -0.156 | 0.024 | [-0.203, -0.110] | <0.001 | -0.156 | 0.024 | [-0.203, -0.108] | <0.001 |
| **$Age_{BAS}$** | -0.133 | 0.040 | [-0.212, -0.054] | 0.001 | -0.134 | 0.039 | [-0.212, -0.057] | 0.001 | -0.150 | 0.041 | [-0.230, -0.071] | <0.001 |
| **Sex** | 0.148 | 0.081 | [-0.010, 0.307] | 0.067 | 0.166 | 0.080 | [0.010, 0.321] | 0.037 | 0.148 | 0.081 | [-0.011, 0.307] | 0.068 |
| **Tau $load_{BAS}$** | -0.241 | 0.041 | [-0.321, -0.161] | <0.001 | -0.217 | 0.057 | [-0.330, -0.104] | <0.001 | -0.093 | 0.055 | [-0.200, 0.014] | 0.088 |
| **Tau $LI_{BAS}$** | 0.083 | 0.042 | [0.001, 0.164] | 0.046 | -0.044 | 0.057 | [-0.156, 0.068] | 0.444 | -0.143 | 0.054 | [-0.250, -0.037] | 0.008 |

| Time × $Age_{BAS}$ | -0.020 | 0.019 | [-0.057, 0.017] | 0.282 | -0.028 | 0.017 | [-0.062, 0.005] | 0.097 | -0.036 | 0.018 | [-0.071, -0.002] | 0.039 |
|---|---|---|---|---|---|---|---|---|---|---|---|---|
| Time × Sex | -0.033 | 0.037 | [-0.106, 0.040] | 0.373 | -0.035 | 0.034 | [-0.101, 0.031] | 0.304 | -0.035 | 0.034 | [-0.102, 0.032] | 0.306 |
| Time × Tau $load_{BAS}$ | -0.117 | 0.019 | [-0.155, -0.080] | <0.001 | -0.163 | 0.025 | [-0.213, -0.114] | <0.001 | -0.077 | 0.025 | [-0.125, -0.029] | 0.002 |
| Time × Tau $LI_{BAS}$ | -0.010 | 0.019 | [-0.047, 0.028] | **0.611** | -0.015 | 0.025 | [-0.065, 0.034] | **0.549** | -0.104 | 0.025 | [-0.153, -0.055] | **<0.001*** |
| LME #3: **mPACC** ~ **time** * ($Age_{BAS}$ + Sex + **Aβ $LI_{BAS}$**) + [time \| participant] | | | | | | | | | | | | |
| **ROI** | **Braak I-II** | | | | **Braak III-IV** | | | | **Braak V-VI** | | | |
| | **β** | **SE** | **95% CI** | **p** | **β** | **SE** | **95% CI** | **p** | **β** | **SE** | **95% CI** | **p** |
| **Intercept** | 0.142 | 0.060 | [0.023, 0.260] | 0.019 | 0.143 | 0.060 | [0.026, 0.260] | 0.017 | 0.145 | 0.060 | [0.027, 0.263] | 0.016 |
| **Time** | -0.161 | 0.029 | [-0.217, -0.105] | <0.001 | -0.162 | 0.029 | [-0.218, -0.106] | <0.001 | -0.161 | 0.029 | [-0.218, -0.105] | <0.001 |
| **$Age_{BAS}$** | -0.117 | 0.043 | [-0.201, -0.033] | 0.006 | -0.123 | 0.042 | [-0.206, -0.040] | 0.004 | -0.120 | 0.042 | [-0.204, -0.037] | 0.004 |
| **Sex** | 0.165 | 0.086 | [-0.003, 0.334] | 0.054 | 0.162 | 0.085 | [-0.005, 0.328] | 0.057 | 0.155 | 0.086 | [-0.013, 0.322] | 0.070 |
| **Aβ $LI_{BAS}$** | -0.045 | 0.043 | [-0.129, 0.039] | 0.297 | 0.116 | 0.043 | [0.033, 0.200] | 0.006 | 0.089 | 0.042 | [0.006, 0.172] | 0.035 |
| **Time × $Age_{BAS}$** | -0.016 | 0.021 | [-0.057, 0.024] | 0.427 | -0.018 | 0.021 | [-0.058, 0.023] | 0.392 | -0.017 | 0.021 | [-0.058, 0.023] | 0.397 |
| **Time × Sex** | -0.019 | 0.041 | [-0.099, 0.061] | 0.639 | -0.019 | 0.041 | [-0.099, 0.061] | 0.645 | -0.020 | 0.041 | [-0.100, 0.060] | 0.626 |
| **Time × Aβ $LI_{BAS}$** | -0.012 | 0.021 | [-0.052, 0.029] | **0.572** | 0.016 | 0.020 | [-0.024, 0.056] | **0.420** | 0.012 | 0.020 | [-0.029, 0.052] | **0.573** |

### Replication in independent cohorts

***Table S2.9**. Demographics of the external cohorts.*
*Categorical variables have been presented as 'count (%)', normally distributed continuous variables as 'mean (SD)' and non-normally distributed variables as 'median [IQR]'. OASIS-3 – Open Access Series of Imaging Studies; A4 – Anti-Amyloid Treatment in Asymptomatic Alzheimer's Disease; ADNI – Alzheimer's Disease Neuroimaging Initiative; M – male; F – female; CU – cognitively unimpaired; CI – cognitively impaired; SUVR – standardised uptake value ratio; Aβ – amyloid-beta.*

| | | **OASIS-3 (A+T+; n=46)** | **A4 (A+T+; n=55)** | **ADNI (A+T+; n=133)** |
|---|---|---|---|---|
| **Age, years** | | 74.61 (6.77) | 72.65 (5.06) | 72.39 (6.72) |
| **Sex** | **M** | 21 (46%) | 21 (38%) | 56 (42%) |
| | **F** | 25 (54%) | 34 (62%) | 77 (58%) |
| **Education, years** | | 15.78 (2.60) | 16.60 (2.66) | 15.69 (2.40) |
| **Diagnosis** | **CU** | 15 (33%) | 55 (100%) | 23 (17%) |
| | **CI** | 31 (67%) | 0 (0%) | 110 (83%) |

| | | | |
|---|---|---|---|
| **Temporal tau, SUVR** | 1.57 [1.45,1.85] | 1.38 [1.34,1.46] | 1.57 [1.39,1.82] |
| **Global Aβ, SUVR** | 1.40 [1.27,1.59] | 1.37 [1.27,1.46] | 1.42 [1.32,1.53] |

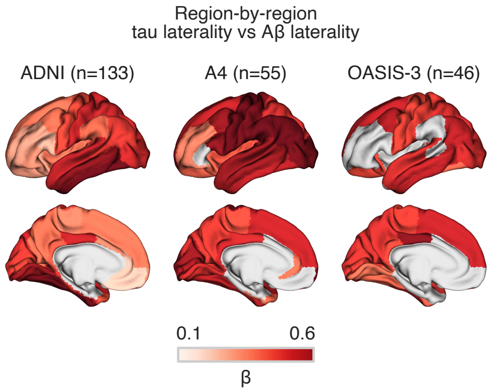


***Figure S2.6***. *Region-by-region associations between Aβ laterality and tau laterality in external cohorts. Aβ – amyloid-beta; ADNI – Alzheimer's Disease Neuroimaging Initiative; A4 – Anti-Amyloid Treatment in Asymptomatic Alzheimer's Disease; OASIS-3 – Open Access Series of Imaging Studies*

### Sensitivity analyses

##### Partial volume corrected PET

We performed cross-sectional and longitudinal sensitivity analyses using partial volume corrected PET SUVR values to investigate if the association between laterality of Aβ and tau distribution could be related to this methodological choice. Cross-sectionally, the analysis showed a similarly strong association between Aβ laterality and tau laterality in all main meta-ROIs (Fig. S2.7; Global: β=0.648, p<0.001; Braak I-II: β=0.178, p=0.005; Braak III-IV: β=0.705, p<0.001; Braak V-VI: β=0.590, p<0.001). One outlier was removed from the analysis based on the 4 SD difference from the mean. Moreover, the longitudinal sensitivity analysis provided similar results to the main results with the full A+ sample where higher baseline Aβ laterality was predictive of changes over time in tau laterality (Fig. S2.8a; β=0.041, 95%CI=[0.021; 0.060], p<0.001). Similarly, A+T- group showed significant interaction effect only in Braak I-II (Fig. S2.8b; β=0.078, 95%CI=[0.027; 0.129], p=0.009) and Braak III-IV (β=0.070, 95%CI=[0.021; 0.119], p=0.015). However, A+T+ group exhibited significant effect on baseline Aβ laterality on tau laterality over time in Braak V-VI (Fig. S2.8c; β=0.064, 95%CI=[0.034; 0.094], p<0.001), which we did not identify in the main findings. For individuals who stayed A+T- throughout their follow-up, we detected a significant interaction effect only in Braak I-II (Fig. S2.8d; β=0.113, 95%CI=[0.050; 0.175], p=0.001), which was in accordance with the main findings, but for people who converted to A+T+ we did not find any significant association in any of the Braak meta-ROIs in contrast to our main findings – only trending level significance in Braak V-VI (Fig. S2.8e; β=0.093, 95%CI=[0.014; 0.171], p=0.061) and no significance in Braak III-IV (β=0.064, 95%CI=[-0.012; 0.141], p=0.302).

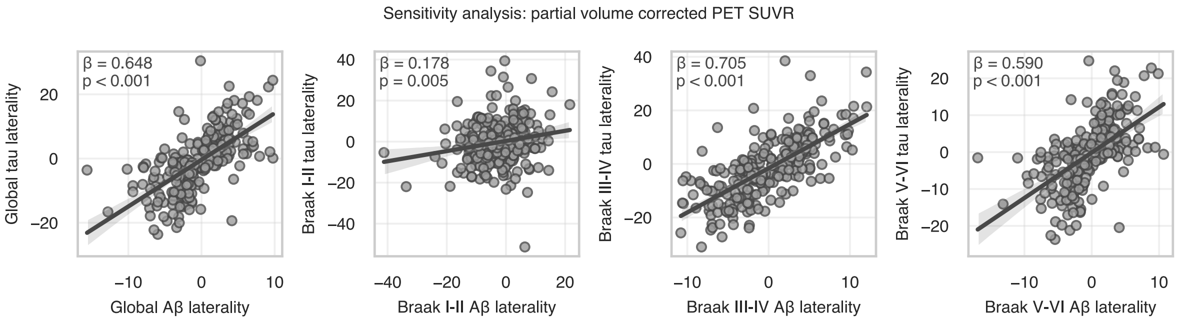


***Figure S2.7****. Cross-sectional association between Aβ and tau laterality with using partial volume corrected SUVR values for calculating laterality.*

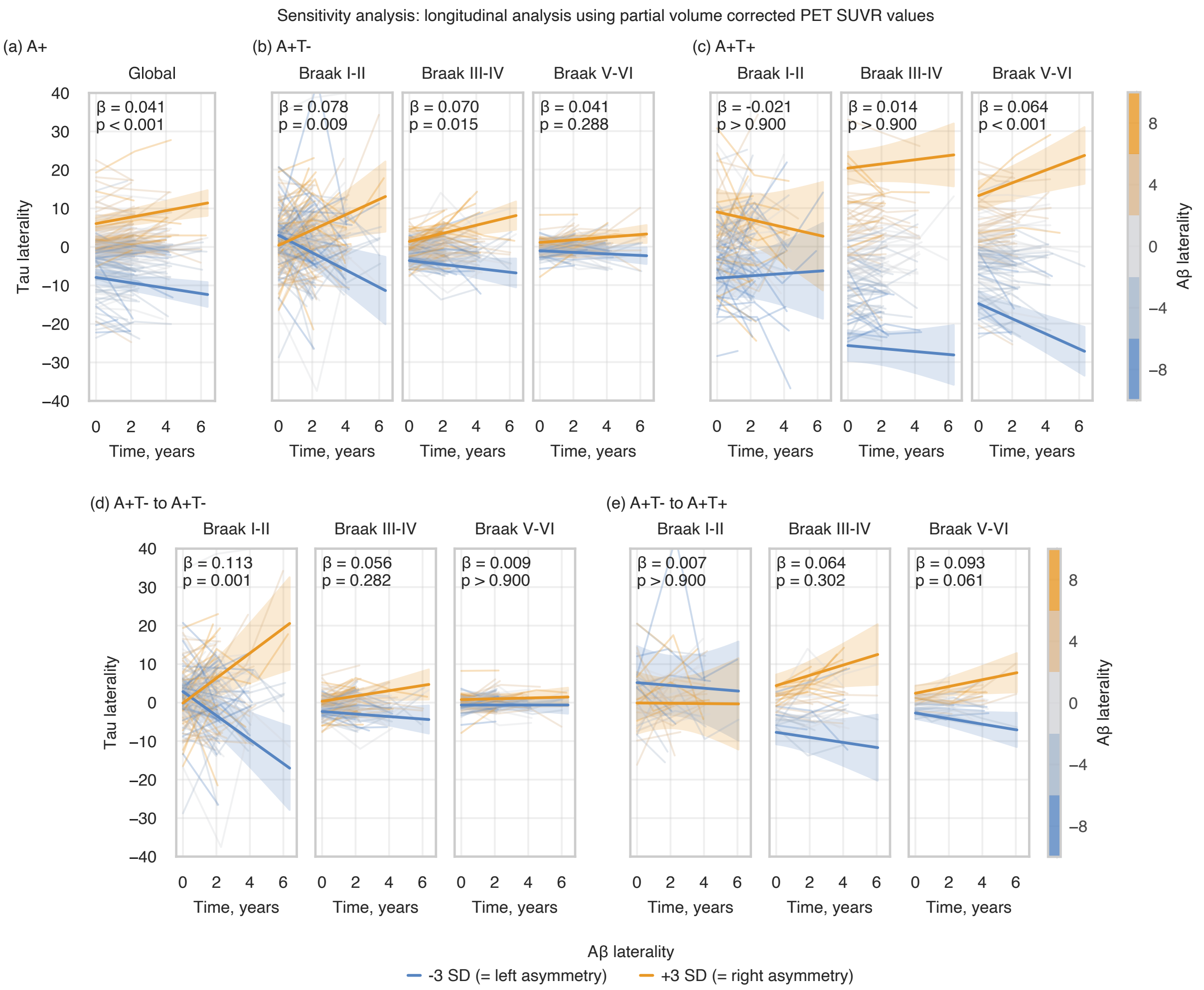


***Figure S2.8***. *Longitudinal analysis of the association between baseline Aβ laterality and changes over time in tau laterality at Braak meta-ROIs using partial volume corrected SUVR value for calculating laterality: (a) whole A+ sample at global meta-ROI (i.e., whole-brain for each hemisphere); (b) A+T- subsample at Braak meta-ROIs; (c) A+T+ subsample at Braak meta-ROIs; (d) A+T- subsample who stay A+T- throughout their follow-up; (e) A+T- subsample who progress to A+T+ during their follow-up.*
*The statistical annotations indicate the effect size and significance level of the interaction between time and baseline Aβ laterality on tau laterality. For visualisation, regression lines with 95% CIs of ±3 SD baseline Aβ laterality were plotted. Aβ – amyloid-beta*

#### Association of the distribution of Aβ and tau with cerebral blood flow and cortical thickness

To test whether our main cross-sectional finding of a strong association between Aβ and tau distribution was not due to other biological confounders, we tested to what extent Aβ laterality and tau laterality are associated to laterality in cerebral blood flow and cortical thickness. Cerebral blood flow was estimated using arterial spin labelling (ASL) scans which were acquired on a subset of the sample and its methodology in detail has been described previously.[35] Cortical thickness was measured as the distance from the grey matter/white matter boundary to the corresponding pial surface.[36] Only tau laterality was negatively related to laterality in cerebral blood flow (n=101; β=-0.527, p<0.001), but not Aβ (n=53; β=-0.241, p=0.085) (Fig. S2.9a). Furthermore, the laterality indexes of both pathologies were negatively associated with laterality in cortical thickness, but with greatly stronger effect with tau (n=449; β=-0.629, p<0.001) than Aβ (n=231; β=-0.185, p=0.005) (Fig. S2.9c). Most importantly, the association between the laterality of Aβ and tau distribution was still statisitically significant after adjusting for the laterality of either cerebral blood flow (Fig. S2.9b; n=53; β=0.498, p<0.001) or cortical thickness (Fig. S2.9d; n=231; β=0.560, p<0.001).

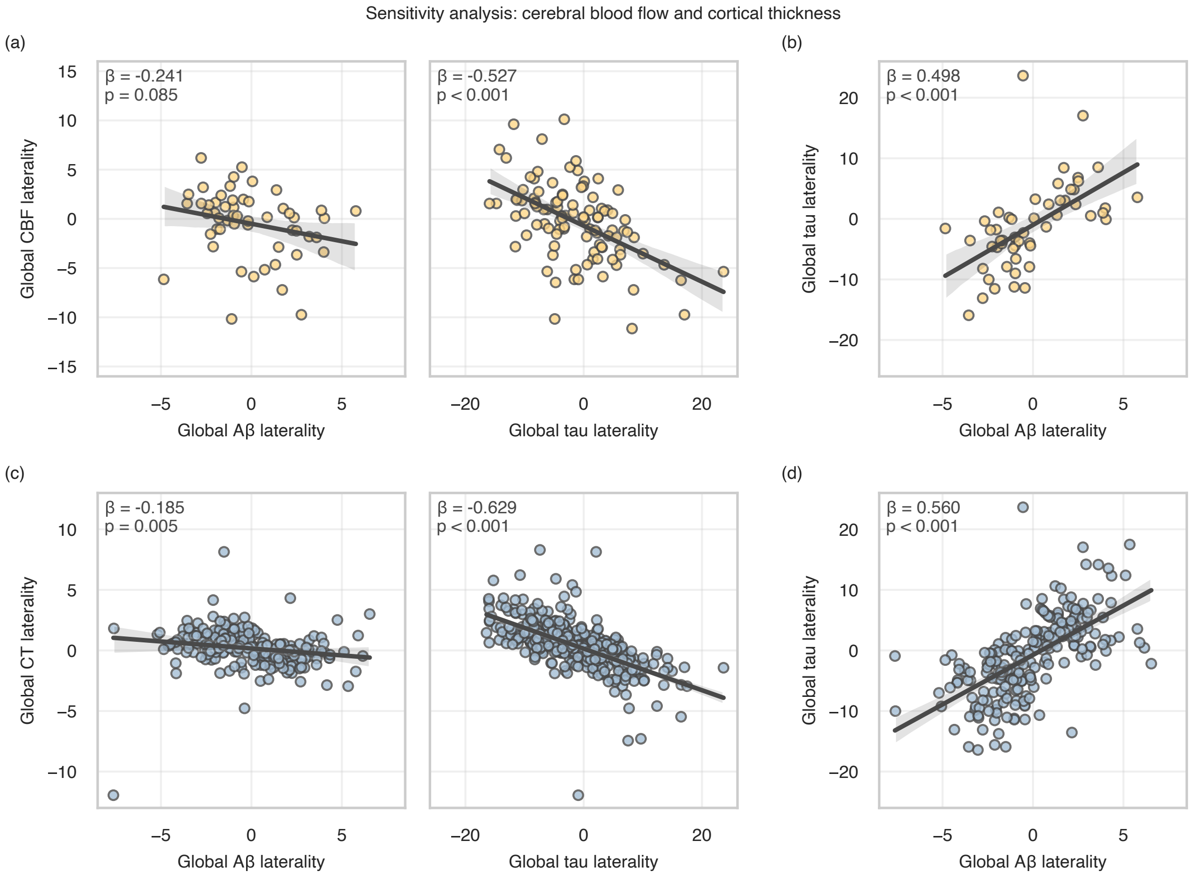


***Figure S2.9**. Cross-sectional associations between the laterality of both Aβ and tau to the laterality of CBF and CT: (a) between the laterality of Aβ/tau and CBF; (b) between the laterality of Aβ and tau after adjusting for CBF; (c) between the laterality of Aβ/tau and CT; (d) between the laterality of Aβ and tau after adjusting for CT.*

*Aβ – amyloid-beta; CBF – cerebral blood flow; CT – cortical thickness*

### Supplementary S3 – PET scans of representative cases

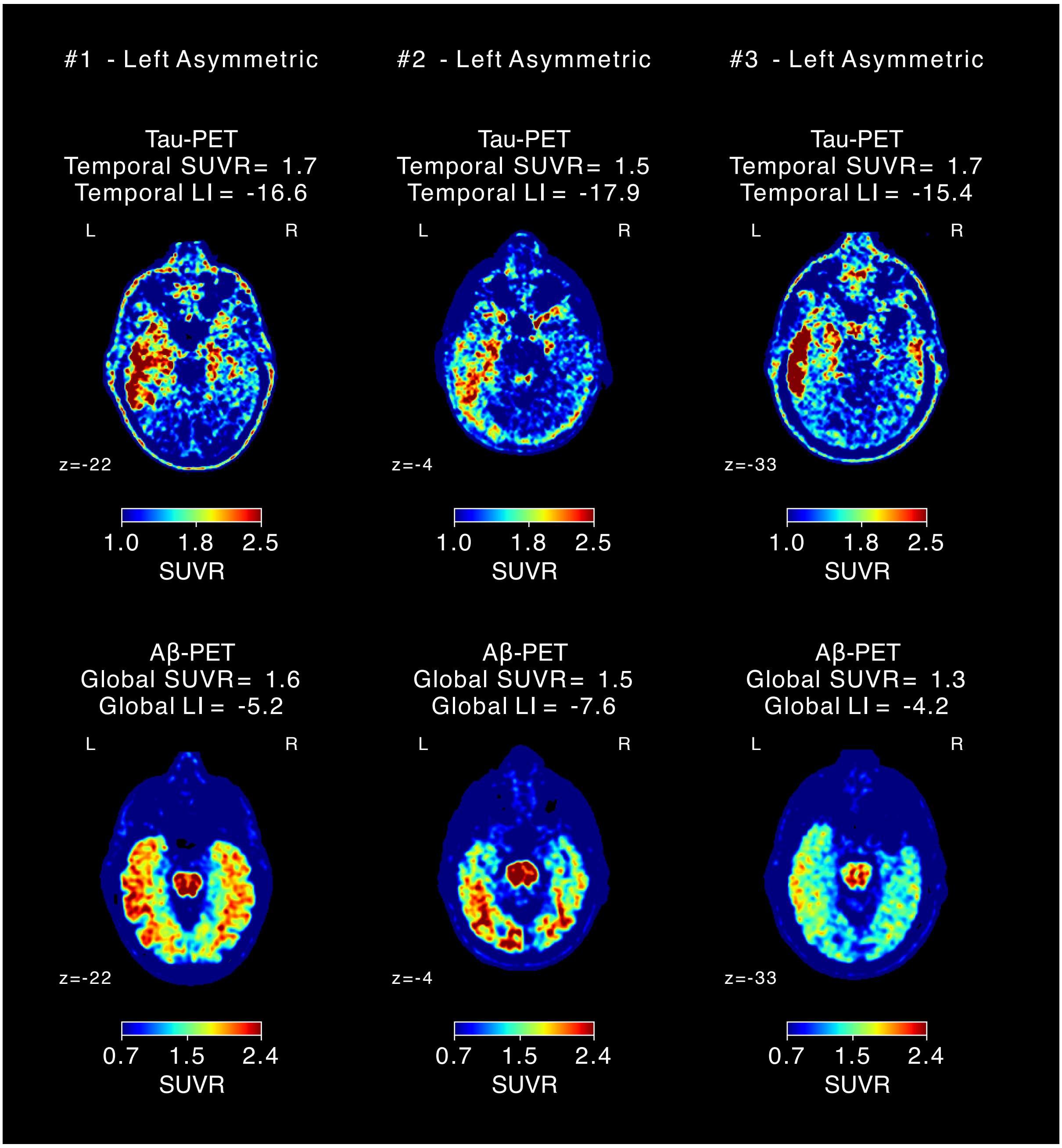


***Figure S3.1***. *Visualisation of tau-PET and Aβ-PET scans for three representative cases with left asymmetric pathological distribution.*
*SUVR – standardised uptake value ratio; LI – laterality index*

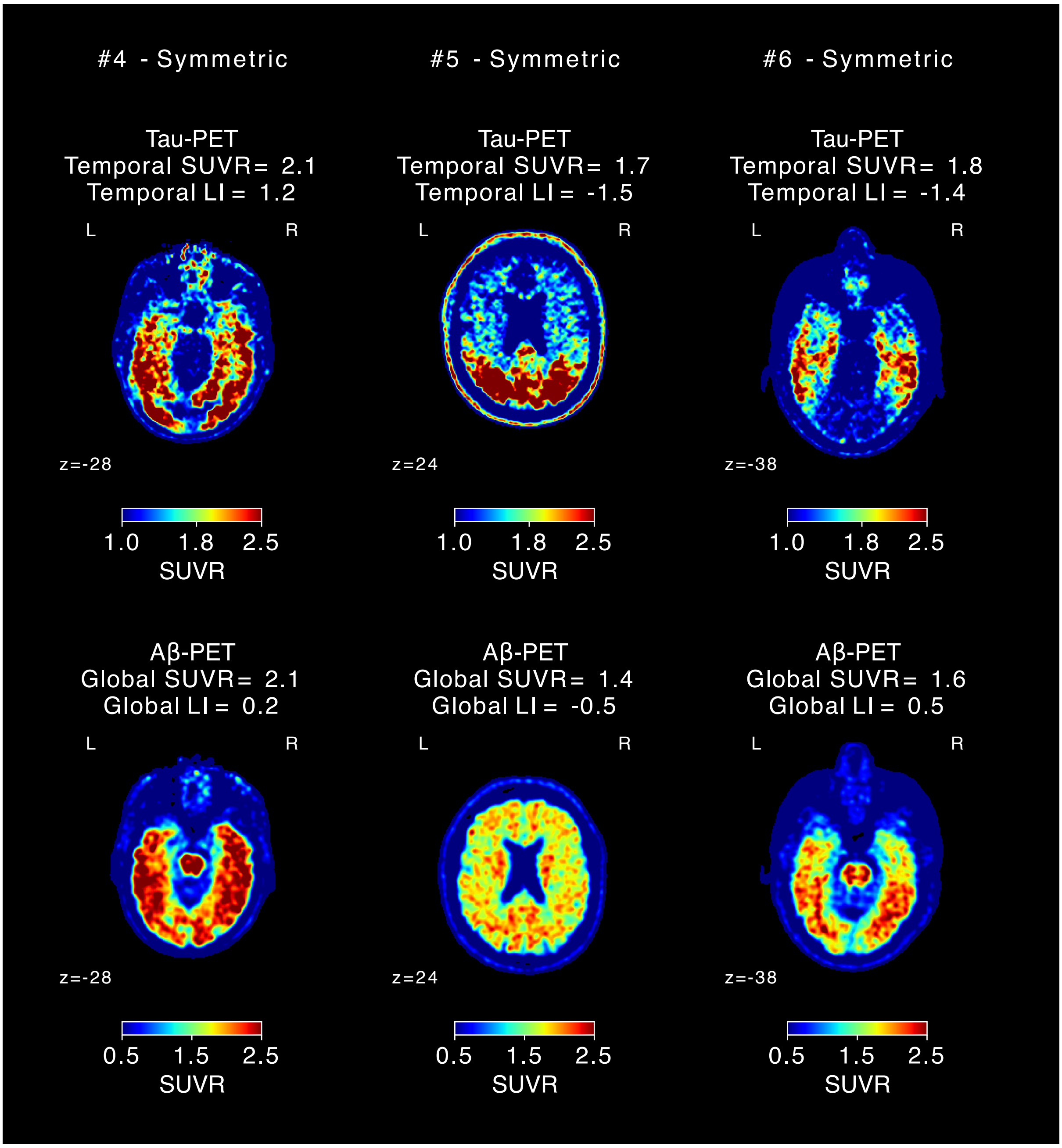


***Figure S3.2***. *Visualisation of tau-PET and Aβ-PET scans for three representative cases with symmetric pathological distribution.*
*SUVR – standardised uptake value ratio; LI – laterality index*

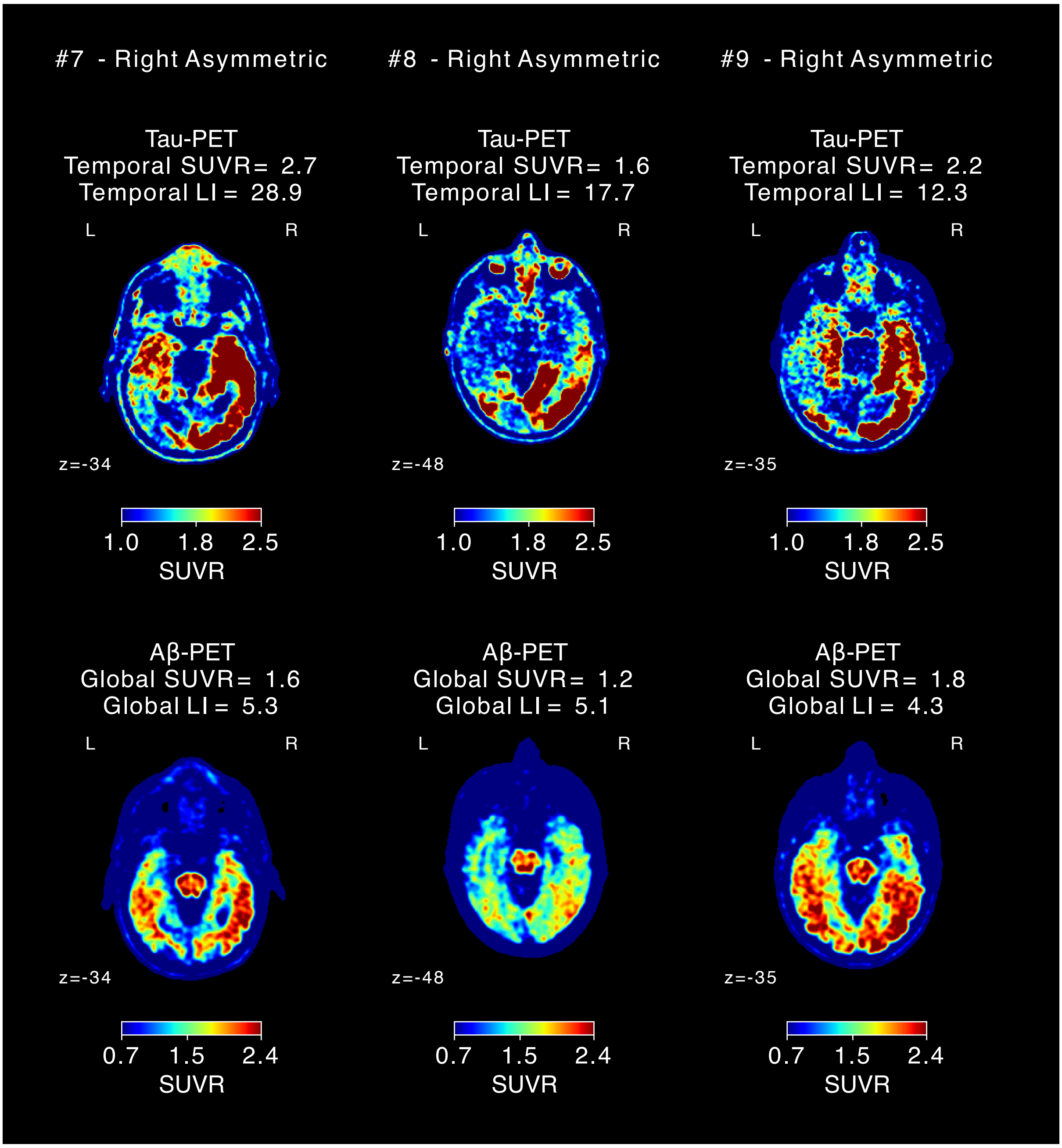


***Figure S3.3***. *Visualisation of tau-PET and Aβ-PET scans for three representative cases with right asymmetric pathological distribution.*
*SUVR – standardised uptake value ratio; LI – laterality index*

### References


1. American Psychiatric Association. *Diagnostic and Statistical Manual of Mental Disorders*. Fifth Edition. American Psychiatric Association; 2013. doi:10.1176/appi.books.9780890425596

2. Palmqvist S, Janelidze S, Quiroz YT, et al. Discriminative Accuracy of Plasma Phospho-tau217 for Alzheimer Disease vs Other Neurodegenerative Disorders. *JAMA*. 2020;324(8):772-781. doi:10.1001/jama.2020.12134

3. Gelb DJ, Oliver E, Gilman S. Diagnostic Criteria for Parkinson Disease. *Arch Neurol*. 1999;56(1):33-39. doi:10.1001/archneur.56.1.33

4. Litvan I, Agid Y, Calne D, et al. Clinical research criteria for the diagnosis of progressive supranuclear palsy (Steele-Richardson-Olszewski syndrome). *Neurology*. 1996;47(1):1-9. doi:10.1212/WNL.47.1.1

5. Gilman S, Wenning GK, Low PA, et al. Second consensus statement on the diagnosis of multiple system atrophy. *Neurology*. 2008;71(9):670-676. doi:10.1212/01.wnl.0000324625.00404.15

6. Armstrong MJ, Litvan I, Lang AE, et al. Criteria for the diagnosis of corticobasal degeneration. *Neurology*. 2013;80(5):496-503. doi:10.1212/WNL.0b013e31827f0fd1

7. Gorno-Tempini ML, Hillis AE, Weintraub S, et al. Classification of primary progressive aphasia and its variants. *Neurology*. 2011;76(11):1006-1014. doi:10.1212/WNL.0b013e31821103e6

8. Braak H, Braak E. Neuropathological stageing of Alzheimer-related changes. *Acta Neuropathol (Berl)*. 1991;82(4):239-259. doi:10.1007/BF00308809

9. Braak H, Alafuzoff I, Arzberger T, Kretzschmar H, Del Tredici K. Staging of Alzheimer disease-associated neurofibrillary pathology using paraffin sections and immunocytochemistry. *Acta Neuropathol (Berl)*. 2006;112(4):389-404. doi:10.1007/s00401-006-0127-z

10. Mattsson N, Palmqvist S, Stomrud E, Vogel J, Hansson O. Staging β-Amyloid Pathology With Amyloid Positron Emission Tomography. *JAMA Neurol*. 2019;76(11):1319-1329. doi:10.1001/jamaneurol.2019.2214

11. Esteban O, Markiewicz CJ, Burns C, et al. nipy/nipype: 1.8.3. Published online July 14, 2022. doi:10.5281/zenodo.6834519

12. Tournier JD, Smith R, Raffelt D, et al. MRtrix3: A fast, flexible and open software framework for medical image processing and visualisation. *NeuroImage*. 2019;202:116137. doi:10.1016/j.neuroimage.2019.116137

13. Jenkinson M, Beckmann CF, Behrens TEJ, Woolrich MW, Smith SM. FSL. *NeuroImage*. 2012;62(2):782-790. doi:10.1016/j.neuroimage.2011.09.015

14. Fischl B. FreeSurfer. *NeuroImage*. 2012;62(2):774-781. doi:10.1016/j.neuroimage.2012.01.021

15. Dhollander T, Raffelt D, Connelly A. Unsupervised 3-tissue response function estimation from single-shell or multi-shell diffusion MR data without a co-registered T1 image. In: *ISMRM Workshop on Breaking the Barriers of Diffusion MRI, 2016, 5.* ; 2016.

16. Dhollander T, Mito R, Raffelt D, Connelly A. Improved white matter response function estimation for 3-tissue constrained spherical deconvolution. In: *Proc Intl Soc Mag Reson Med, 2019, 555.* ; 2019.

17. Tournier JD, Calamante F, Gadian DG, Connelly A. Direct estimation of the fiber orientation density function from diffusion-weighted MRI data using spherical deconvolution. *NeuroImage*. 2004;23(3):1176-1185. doi:10.1016/j.neuroimage.2004.07.037

18. Jeurissen B, Tournier JD, Dhollander T, Connelly A, Sijbers J. Multi-tissue constrained spherical deconvolution for improved analysis of multi-shell diffusion MRI data. *NeuroImage*. 2014;103:411-426. doi:10.1016/j.neuroimage.2014.07.061

19. Smith R, Skoch A, Bajada C, Caspers S, Connelly A. Hybrid Surface-Volume Segmentation for improved Anatomically-Constrained Tractography. In: *Proceedings of the Oganisation for Human Brain Mapping*. ; 2020.

20. Jenkinson M, Smith S. A global optimisation method for robust affine registration of brain images. *Med Image Anal*. 2001;5(2):143-156. doi:10.1016/s1361-8415(01)00036-6

21. Jenkinson M, Bannister P, Brady M, Smith S. Improved optimization for the robust and accurate linear registration and motion correction of brain images. *NeuroImage*. 2002;17(2):825-841. doi:10.1016/s1053-8119(02)91132-8

22. Tournier JD, Calamante F, Connelly A. Improved probabilistic streamlines tractography by 2nd order integration over fibre orientation distributions. In: *Proceedings of the International Society for Magnetic Resonance in Medicine, 2010, 1670.* ; 2010.

23. Smith RE, Tournier JD, Calamante F, Connelly A. Anatomically-constrained tractography: Improved diffusion MRI streamlines tractography through effective use of anatomical information. *NeuroImage*. 2012;62(3):1924-1938. doi:10.1016/j.neuroimage.2012.06.005

24. Smith RE, Tournier JD, Calamante F, Connelly A. SIFT2: Enabling dense quantitative assessment of brain white matter connectivity using streamlines tractography. *NeuroImage*. 2015;119:338-351. doi:10.1016/j.neuroimage.2015.06.092

25. Desikan RS, Ségonne F, Fischl B, et al. An automated labeling system for subdividing the human cerebral cortex on MRI scans into gyral based regions of interest. *NeuroImage*. 2006;31(3):968-980. doi:10.1016/j.neuroimage.2006.01.021

26. Fischl B, Salat DH, Busa E, et al. Whole Brain Segmentation: Automated Labeling of Neuroanatomical Structures in the Human Brain. *Neuron*. 2002;33(3):341-355. doi:10.1016/S0896-6273(02)00569-X

27. Smith RE, Tournier JD, Calamante F, Connelly A. The effects of SIFT on the reproducibility and biological accuracy of the structural connectome. *NeuroImage*. 2015;104:253-265. doi:10.1016/j.neuroimage.2014.10.004

28. Basser PJ, Mattiello J, Lebihan D. Estimation of the Effective Self-Diffusion *Tensor* from the NMR Spin Echo. *J Magn Reson B*. 1994;103(3):247-254. doi:10.1006/jmrb.1994.1037

29. Basser PJ, Mattiello J, LeBihan D. MR diffusion tensor spectroscopy and imaging. *Biophys J*. 1994;66(1):259-267. doi:10.1016/S0006-3495(94)80775-1

30. Wasserthal J, Neher P, Maier-Hein KH. TractSeg - Fast and accurate white matter tract segmentation. *NeuroImage*. 2018;183:239-253. doi:10.1016/j.neuroimage.2018.07.070

31. Abraham A, Pedregosa F, Eickenberg M, et al. Machine learning for neuroimaging with scikit-learn. *Front Neuroinformatics*. 2014;8. doi:10.3389/fninf.2014.00014

32. Friston KJ. Functional and effective connectivity: a review. *Brain Connect*. 2011;1(1):13-36. doi:10.1089/brain.2011.0008

33. Fisher RA. On the" probable error" of a coefficient of correlation deduced from a small sample. Published online 1921.

34. Betthauser TJ, Bilgel M, Koscik RL, et al. Multi-method investigation of factors influencing amyloid onset and impairment in three cohorts. *Brain*. 2022;145(11):4065-4079. doi:10.1093/brain/awac213

35. Ahmadi K, Pereira JB, Berron D, et al. Gray matter hypoperfusion is a late pathological event in the course of Alzheimer's disease. *J Cereb Blood Flow Metab Off J Int Soc Cereb Blood Flow Metab*. 2023;43(4):565-580. doi:10.1177/0271678X221141139

36. Fischl B, Dale AM. Measuring the thickness of the human cerebral cortex from magnetic resonance images. *Proc Natl Acad Sci*. 2000;97(20):11050-11055. doi:10.1073/pnas.200033797
